# Historical use of coastal wetlands by small-scale fisheries in the Northern Gulf of California

**DOI:** 10.1101/2022.10.31.513536

**Authors:** Hem Nalini Morzaria-Luna, Mabilia Urquidi, Gabriela Cruz-Piñón, Jośe Manuel Dorantes Hernández, Paloma A. Valdivia Jiménez, Ángeles Sánchez Cruz, Ilse Martínez

## Abstract

Coastal wetlands are rich and productive ecosystems that historically have been used by small-scale fisheries due to their role as refuges, feeding, and nursery habitats for commercial target species. We used wetland resource users’ Local Ecological Knowledge to document historical patterns of commercial species abundance, areas of fishing importance, trophic level, and species richness and composition in coastal wetlands in the Northern Gulf of California, Mexico. We also reconstructed the environmental history of coastal wetlands in this region from bibliographic sources and photographic records, to document impacts that could have affected coastal fisheries. We found a consistent downward trend in target species abundance; the decrease was perceived as more pronounced by fishers that began fishing in or prior to the 1950’s, pointing to shifting baselines, the failure for resource users to recognize environmental change and accept degraded states as normal. Areas of fishing importance within coastal wetlands also decreased through time. Trophic level of catch showed no distinct pattern across wetland sites or time. Perceived species richness and composition increased with wetland size. Our analysis of the small-scale use of coastal wetlands in the Northern Gulf is relevant to food security and can provide insight into how local populations adapt to depleted coastal food webs.

## Introduction

Small-scale fisheries (also called artisanal), are defined by labor intensive strategies, low catch capacity, small boats (≤10–15m), short travel distances, a small crew, and limited capital investment (Cánovas-Molina et al. 2021); they represent more than 99% of the world’s 51 million fishers and account for more than half of the total global fisheries production (Jones et al. 2018). These fisheries also play a key social and economic role in coastal communities (Basurto et al. 2013); thus, the engagement of small-scale fishers is a requisite for successful environmental policy, and may influence outcomes more than the policies themselves (Gutiérrez et al. 2011). However, social and political history of a place and fishers’ context must be recognized and understood before enacting policies (Cisneros-Montemayor and Vincent 2016).

Historically, coastal wetlands have been preferred as sites for small-scale fisheries due to their rich and productive ecosystems, and their role as refuges, feeding, and nursery habitats for commercial species (Glenn et al. 2006; Spackeen 2009; Alvirde-López 2012; Turk-Boyer et al. 2014b). In general, coastal wetlands, along with other coastal ecosystems, have suffered major structural and functional changes over the last several centuries as a result of the number and size of target species removed and the spatial distribution of fishing activities (Sáenz-Arroyo et al. 2005a; Lotze et al. 2006; Kennett et al. 2008; Lotze 2010; Beaudreau and Whitney 2016). These changes include decreases in species abundance and biomass, particularly of large vertebrates and shellfish (Ainsworth 2011; Rick et al. 2016) and are evident in the reduction of the mean trophic level of fisheries landings since the start of industrialized fishing, in what has been termed as “fishing down food webs” (Pauly et al. 1998a) and in the extinction or collapse of target and bycatch species (Jackson et al. 2001). Overfishing interacts with other impacts, such as pollution, sediment and nutrient loading, habitat destruction, and climate change, leading to synergistic responses and threshold effects (Jackson et al. 2001; Scheffer et al. 2005).

Documenting the effects of fishing on wetland ecosystems is important for ecological modeling, management, restoration, and conservation (Jackson et al. 2001; McClenachan et al. 2012), particularly because the true magnitude of declines can be masked by ‘shifting baselines’ (Pauly 1995), where each consecutive generation accepts a lower standard of resource abundance as being normal (Ainsworth et al. 2008a). This failure to recognize the extent of past environmental modifications by humans can apply to both resource users (Sáenz-Arroyo et al. 2005a) and scientists alike (Baum and Myers 2004).

Documenting ‘historical’ use of coastal wetlands by small-scale fisheries can be achieved by integrating multiple knowledge sources, such as oral histories, contemporary fisheries data, historical documentation, and archaeological information (Van Dyke and Wasson 2005; Ames 2007; Ainsworth et al. 2008a; Beaudreau and Levin 2013). Unconventional data sources, such as Local Ecological Knowledge (LEK), the empirical knowledge of a particular group of people that reflects their interactions with local ecosystems and naturalists accounts (Sáenz-Arroyo et al. 2005a) can help reconstruct species abundance trends (Olsson and Folke 2001; Lozano-Montes et al. 2008), species declines (Sáenz-Arroyo et al. 2005b; Turvey et al. 2010; Azzurro et al. 2011) and to understand local ecological processes (Silvano and Valbo-Jørgensen 2008).

LEK has proven to be a cost-effective way to collect high-quality ecological data (Leduc et al. 2021). In data-poor systems, such as the Gulf of California, LEK may be the only available source of historical information to document long-term exploitation of coastal fishery resources (Robertson and McGee 2003; Beaudreau and Levin 2013; Rubio-Cisneros et al. 2017).

Here, we document historical patterns of commercial species abundance, areas of fishing importance, trophic level, and species richness and composition in coastal wetlands in the Northern Gulf of California, Mexico. The analysis of the patterns of small-scale fishing in coastal wetlands is relevant to food security and can provide insight into how local populations adapt to depleted coastal food webs (Rubio-Cisneros et al. 2017). The Gulf of California is a highly productive system that supports the most economically important fisheries in Mexico (Cisneros-Mata 2010). Most coastal communities in the Gulf of California depend on a small-scale fishing sector that targets over 80 species of fish, sharks, and invertebrates (Turk-Boyer et al. 2014a); these fisheries are highly adaptive and flexible, and their multispecific and multigear character allows fishers to maximize profitability (Arce-Acosta et al. 2018). Changes in the marine environment of the Gulf of California, including decreases in catch composition and mean trophic level (Sala et al. 2004), maximum fish size (Sala et al. 2004; Sáenz-Arroyo et al. 2005b), and species abundance (Sáenz-Arroyo et al. 2005a, b; Lozano-Montes et al. 2008; Ainsworth 2011), have been linked to overfishing.

We elicited the LEK of resource users through oral histories, participatory mapping, and participatory drawing to characterize species abundance trends, important fishing areas, changes in the trophic level of catch between between the 1950’s - 1990’s. These approaches have proven useful to document artisanal fisheries areas and effort (Moreno-Báez et al. 2012; Previero and Gasalla 2018; Chan et al. 2019), changes in the spatial distribution of fisheries over time (Coll and Lotze 2016; Frans and Augé 2016), abundance trends (Robertson and McGee 2003; Ainsworth et al. 2008b; Ainsworth 2011), and essential fish habitats (Serra-Pereira et al. 2014; DeCelles et al. 2017). We also reconstructed the environmental history of coastal wetlands in the Northern Gulf from bibliographic sources and photographic records, to document possible habitat changes that could have affected coastal fisheries.

## Methods

We documented abundance trends, areas of fishing importance, trophic level, and richness and composition of target species extracted by small-scale fisheries in coastal wetlands in the Northern Gulf of California along the Puerto Peñasco -Puerto Lobos Coastal Corridor, Sonora, Mexico (Figure 1). We combined the Local Ecological Knowledge (LEK) of wetland resource users (i.e. small-scale fishers and oyster farmers) with expert interviews to reconstruct historical ecosystem trends in species abundance for commercial species, fishing areas, species richness, and composition in coastal wetlands. Modern occupation of the Northern Gulf did not begin until the 1930’s with the settlement of the first fishing villages and the construction of the railway that connected the region to the rest of Mexico (Enríquez-Acosta 2008). Thus, we were still able to record the LEK of resource users that arrived with the first wave of modern occupation.

**Figure 1.**
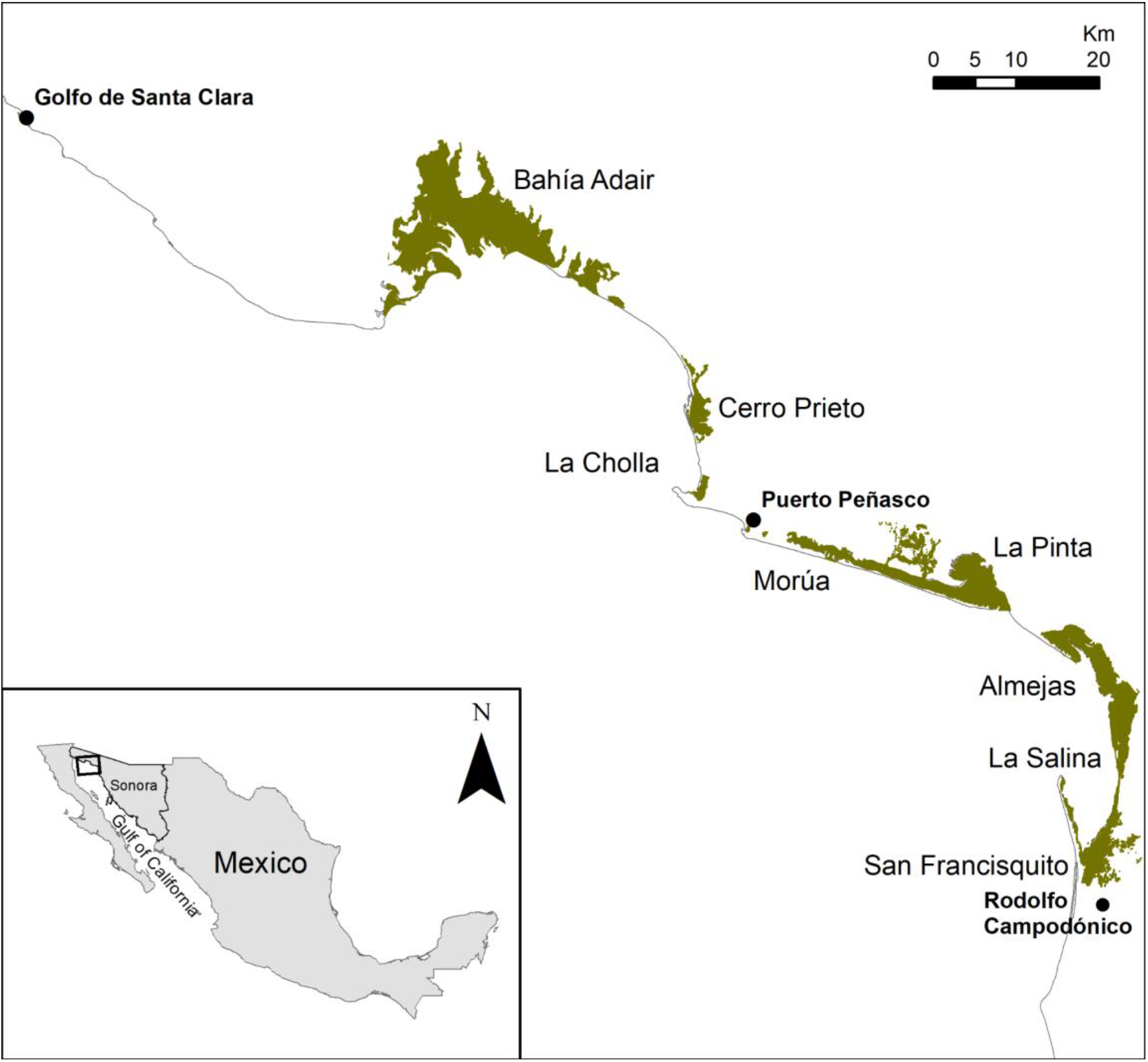
Wetlands in the Northern Gulf of California along the Puerto Peñasco Coastal Corridor in Sonora, México. Data: Sierra and Chamberlain (1999). For interpretation of the references to color in this figure legend, the reader is referred to the online version of the article.

We documented oral histories from wetland resource users through interviews that included participatory mapping, participatory drawing, and questions regarding species’ abundance. We then estimated abundance trends for commercial fishery species and changes in areas of fishing importance indices using a fuzzy logic expert system that accounts for the inherent uncertainty in users’ interview responses. We also recorded the environmental history of coastal wetlands in the Northern Gulf to understand drivers of changes in species’ abundance, richness, composition, and fishing areas. The analysis was conducted using the R statistical framework (R Development Core Team 2021) with libraries Vegan (ver. 1.17-4 (Oksanen et al. 2010) and MASS (Venables and Ripley 2002) and ArcMap (ESRI, Redmond, CA).

### a. Study area

Coastal wetlands in the Northern Gulf of California cover a total of 114,206 ha; they are a complex of intertidal mudflats, tidal channels, salt marshes, salt flats, coastal dunes, beaches, and the wetland-terrestrial ecotone (Brusca et al. 2006; Glenn et al. 2006). The wetlands lack perennial freshwater, which combined with the high evaporation rates and a macrotidal regime (tidal amplitude can reach 8m), results in higher salinity at the head than at the mouth (i.e. inverse or negative hypersaline estuaries) (Thomson et al. 1968). In the Northern Gulf, wetlands support a diverse, marine-driven food web (Spackeen 2009), and serve as feeding, resting and nursing areas for a variety of invertebrates, birds, fish and large marine vertebrates, such as sea turtles and dolphins (Calderon-Aguilera et al. 2003; Seminoff and Nichols 2006; Mellink and Orozco-Meyer 2006; Gomez-Sapiens et al. 2013). Seventeen key target species (Table 1) are extracted from these wetlands using gill nets, traps, and cast nets (Cudney-Bueno and Turk Boyer 1998; Moreno-Báez et al. 2012); trawls are banned inside wetlands (DOF 2013) and the depth and extreme tidal regime within these systems limit the use of other gears.

**Table. 1.**
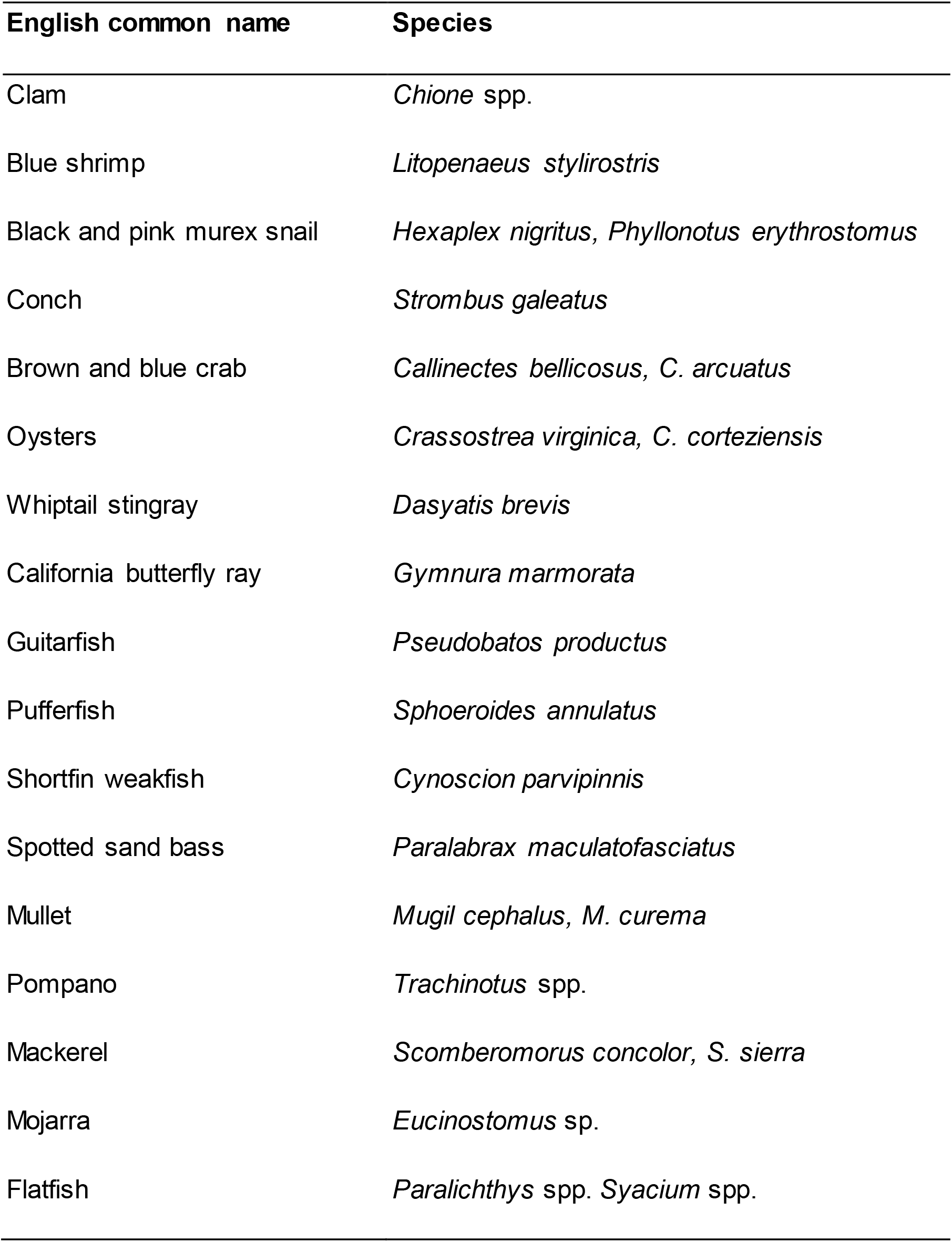
Key species surveyed for abundance trends and fishing areas in fisher interviews (See Supplementary information for Spanish names)

### b. Oral interviews

We carried out the interviews between April and May 2008. Interviewees were selected using snowball sampling, where current subjects recruit their acquaintances to be future subjects (Goodman 1961). The interviews were digitally recorded. Each participant was presented with a written consent form prior to the interview. Since Mexico has no overarching Human-subjects research regulations and participating organizations did not have an Institutional Review Board (IRB), we developed a written consent form following recommendations by The University of Arizona IRB (Texts S1 and S2), that was presented to each interviewee.

We interviewed 48 resource users, using semi-structured oral interviews and participatory mapping and drawing. Resource users lived in the three major communities of the Northern Gulf of California located within the Coastal Corridor, Golfo de Santa Clara (n = 6 interviews), Puerto Peñasco (n = 25), and Ejido Rodolfo Campodónico (n = 17). We developed separate questionnaires for two major categories of resource users, oyster farmers (Text S3) and fishers (Text S4). The information requested in the questionnaire included demographic characteristics, information on species trends, and fishing activities within each wetland, including if resource users had observed changes in breeding areas and extreme reductions in population size. We focused on the 17 invertebrate and fish species most commonly fished in wetlands of the Northern Gulf (Table 1), which were selected based on conversations with expert fishers and on Cudney-Bueno and Turk Boyer (1998). These priority commercial species include wetland dependent species, defined as those that use wetlands for all or part of their life cycle, and non-dependent species. Illustrations of each species were used to verify correspondence between local names and scientific names.

We also carried out eight expert phone interviews. These experts belonged to Universities and non-profit organizations and on average had worked in the region for ~40 years. The questions included an assessment of observed impacts and changes (Text S5).

### c. Species abundance

To estimate temporal shifts in the relative abundance of target species, resource users were asked to describe the relative abundance of each species within five time periods, 1950’s and prior, 1960’s, 1970’s, 1980’s, 1990’s, and 2000’s. We asked resource users to assign to each species an index of change in abundance, as increasing (+1), decreasing (−1), or stable (0), for each of these five periods. Each numeric index was converted into an interview score of high, medium, or low abundance based on a running sum where the number of responses was divided between high, medium and low linguistic scores. We then used a fuzzy expert system (‘Fuzzy logic’; (Zadeh 1965) to combine fishers’ perceptions of species abundance with other population and habitat indicators for each species and time period, according to the methodology developed by Ainsworth et al. (2008b). This method allows integrating linguistic abundance scores with other qualitative and quantitative information in the form of heuristic rules. Fuzzy set theory (or ‘fuzzy logic’) is useful for handling the uncertainty inherent to interview responses, i.e. it is difficult to provide a clear definition of what a response of ‘high abundance’ means, different respondents may have different perceptions or these might change between species. For details on the calculations involved in the fuzzy sets and heuristic rules see Ainsworth et al. (2008b).

#### Wetland habitat Index

We estimated a Wetland habitat Index that indicates the degree of change in each wetland. We calculated the index for each decade between 1950-2000 based on the total wetland area converted to other uses, the proportion of wetlands lost, and the average proportion of expert users that indicated a loss of wetland functions. The total area of wetland habitat lost was obtained from (Sierra and Chamberlain 1999; Glenn et al. 2006; Turk-Boyer 2008). As a proxy for degradation that might not be evident just by the amount of wetland converted to other uses, we used the increase in road construction within wetlands between 1985 and 1997 (Sierra and Chamberlain 1999) and the changes in wetland function derived from the expert interviews. Expert users rated seven wetland functions as increasing, decreasing, or no change; functions could be rated for all habitats or for individual wetlands. We obtained the proportion of respondents that scored a function as decreasing in a time period and estimated the average across all functions, and we weighed the functions directly related to fishing or habitat value.

#### Fuzzification

The average abundance scores, wetland habitat index, changes in breeding areas, and extreme reductions in population size were categorized into one or more levels simultaneously, with the degree of membership to the levels being defined by fuzzy membership functions. The average abundance scores, for each species and decade, were assigned a degree of membership in linguistic categories (Fuzzification), as defined by trapezoid and triangular fuzzy membership functions (Figure 2) (Friedman and Kandel 1999). For instance, based on the membership functions presented in Figure S1, an abundance score of 0.8 can be classified as medium and high abundance with a degree of membership of 0.2 and 0.8, respectively (maximum of 1). We report membership functions weighted between a membership function assuming low variance (high agreement between abundance scores) and high variance (low agreement). We also defined membership functions (as high/low, yes/no) for other qualitative and quantitative indicators: changes in breeding areas, wetland habitat index, and extreme reductions in population size.

**Figure 2.**
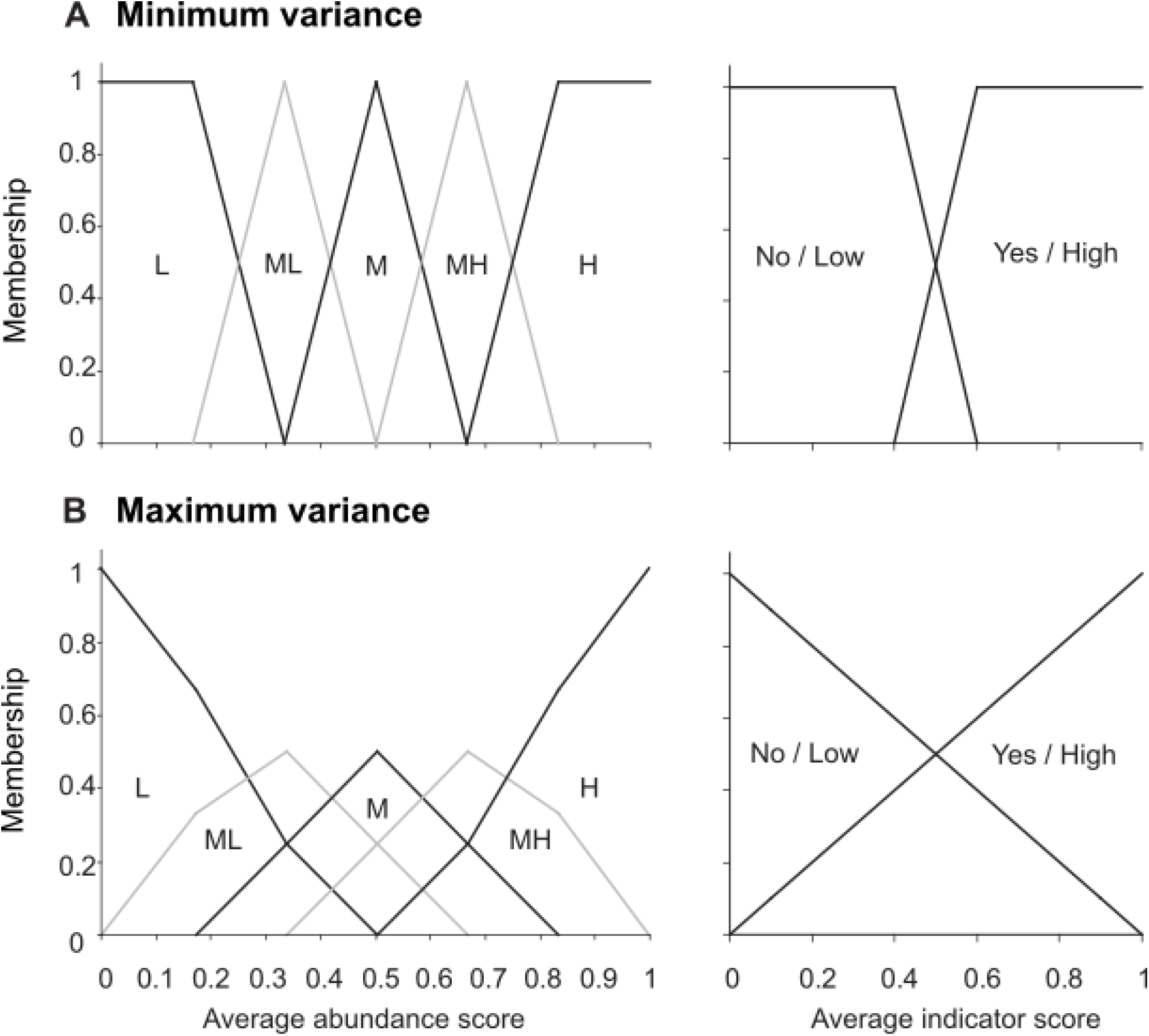
Fuzzy set membership functions. When there is agreement among responses (a.) memberships in the low/yes and high/no categories are defined by trapezoidal functions; memberships in medium low, medium, and medium high categories are defined by triangular functions. As variance among responses increases (b.), the subtended angle increases from a minimum (left) to a maximum (right). X-axes are average abundance scores (left column) or population indicator scores (right column) from interviews for time period and species. The Yes/No membership function was used for the ‘depletion’, ‘breeding’ and ‘habitat’ indicators. Membership categories are L = low, ML = medium low, M = medium, MH = medium-high, H = high.

**Figure 3.**
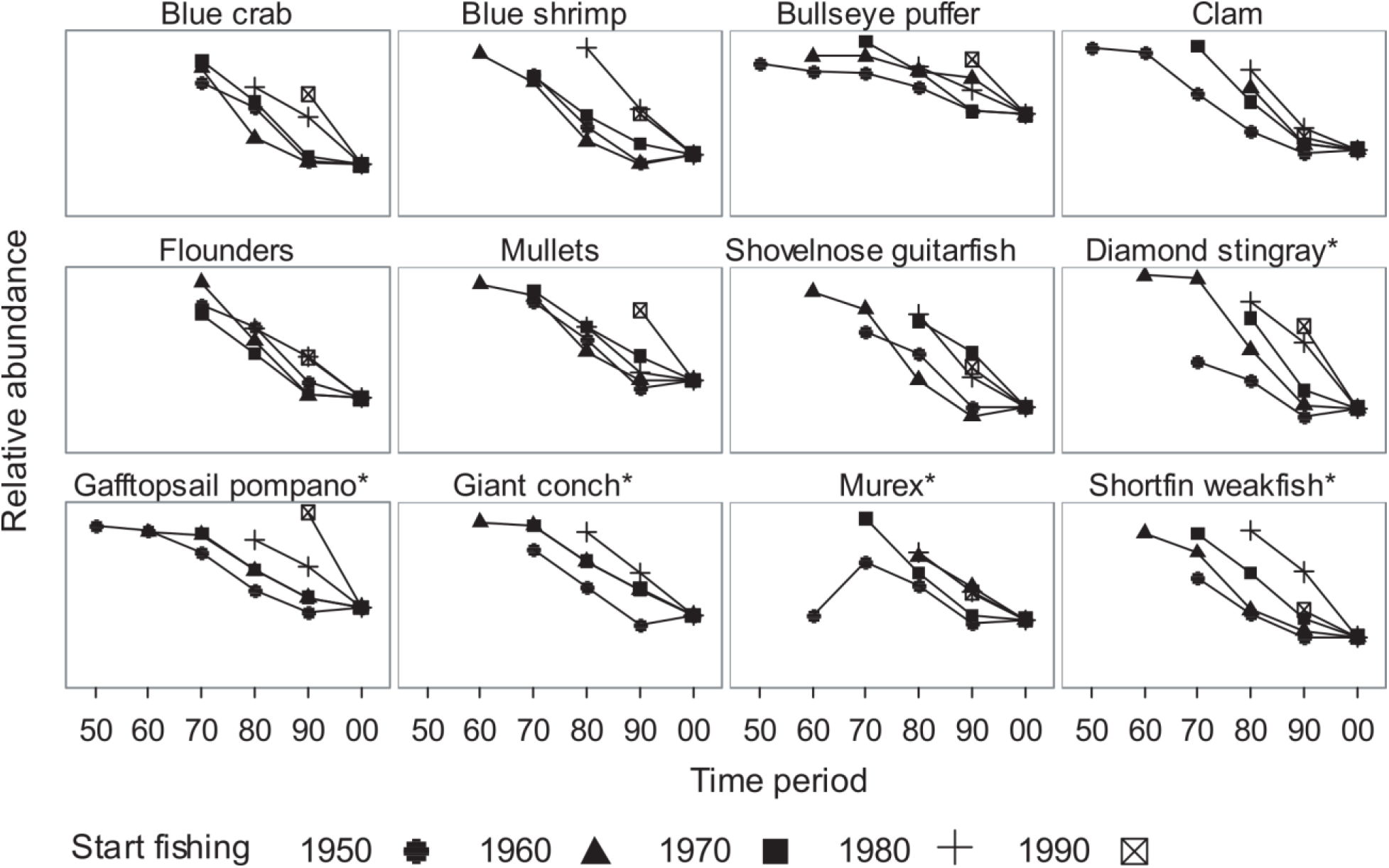
Abundance of 17 target species predicted by the fuzzy expert system, defuzzified using the centroid-weighted average method, based on wetland resource users’ interviews. Values have been standardized to agree in the year 2000. Symbols indicate the decade when the resource users interviewed first reported fishing.

#### Defuzzification

We created heuristic rules of the form: IF a certain situation occurs, THEN a certain outcome is likely. The heuristic rules are captured in a decision table (Table 1S). These rules capture the knowledge contained in the linguistic responses given by interviewees and the qualitative and quantitative indicators. For example, IF wetland area is reduced THEN abundance of wetland-dependent species decreases. Depending on the linguistic variables derived from the membership functions, the rules will fire with a certain strength that reflects the level of belief associated with the rule. We assumed that the rules were triggered only when the abundance score >0. Finally, we reduced the range of conclusions from the different membership functions (Figure 2) into a single output (*Defuzzification*). We obtained the degree of membership of the final conclusion by combining the results of each heuristic rule using an output function of 10 categories to provide a relative index of abundance. We present this index by decade separately for wetland and non-wetland dependent species, defined as those that use the wetland as a nursery and breeding site instead of just for feeding.

### c. Areas of fishing importance

We used participatory mapping to elicit areas of fishing importance from the resource users interviewed. Three interviewees did not complete the participatory mapping portion of the interview, two participants were partially blind and one stated he did not work inside the wetlands. We followed mapping methods from previous studies that used local fishers to identify fishing locations (Close and Brent Hall 2006; Hall and Close 2007). We asked participants to draw the fishing areas for every target species they had knowledge of on maps depicting each coastal wetland; maps varied in scale according to the size of the wetland with a 500 × 500 m grid overlay, except for Bahia Adair which had a 1000 × 1000 grid due to its extent.

We digitized the paper maps using ArcMap 10.3 (ESRI, Redlands, CA) with fishing areas translated into separate polygons. The species and interview information were coded into the feature attribute table. We assigned weights to each polygon based on the number of reported fishing areas by each interviewee (1/number of polygons). We obtained relative weightings, as indicators of a particular area’s importance for fisheries, by summing the values of overlaid individual polygons.

We created vector grid maps corresponding to the printed maps used by the interviewees (500 × 500 m cells for all wetlands except Bahia Adair, which had 1000 × 1000 m cells). Each cell was then used as a standard unit of spatial analysis. We transferred the relative weightings of areas of fishing importance to the vector grid using the weighted average option in the Transfer attributes tool of ET Geowizards (ET Spatial Techniques, Pretoria, South Africa), such that:

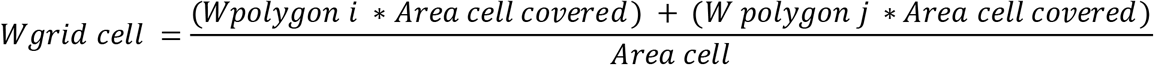

Where *W* is the weight of a given grid cell or polygon.

We used fuzzy logic to estimate historical changes in fishing areas for commercial species by combining fishers’ expert knowledge of where species were fished within each coastal wetland. We grouped interviews based on the decade when the interviewee started fishing in order to compare decades: 1950 and prior, 1960, 1970, 1980, and 1990 and after. Each vector grid was transformed into a raster grid format, constituting the ‘crisp’ sets. We normalized each crisp raster using a fuzzy membership linear function using the Spatial Data Modeler toolbox (Sawatzky et al. 2009) to incorporate uncertainty inherent to interview responses as described by Lanz et al. (2008). Membership in the fuzzy set was expressed in a continuous scale from 1 (full membership) to 0 (full non-membership). This fuzzification method calculates fuzzy membership for a linear transformation of the input crisp raster. The minimum crisp value in each raster was transformed to a membership of 0 and the maximum crisp value was assigned a membership of 1. The membership function then reflects the degree of fishing importance assigned to each wetland by the interviewees. A fuzzy sum operator (Bonham-Carter 1994) was used to combine the fuzzy membership function values of areas indicated by distinct fishers by decade for each wetland. The fuzzy sum operator is equal to

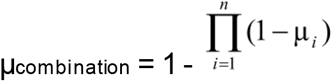

where μ_*i*_ is the fuzzy membership function for the i-th map, and *i*= 1,2,…,n maps are to be combined. This operator has an increasing effect, the result will always be larger (or equal) to the largest fuzzy membership value. In this way two expert judgments of high fishing importance will reinforce one another and the combined judgment is more supportive than either taken individually, although limited by the maximum value of 1.0.

We compared spatial layers of fishing importance using the Kappa statistic (Cohen 1960), which is the most commonly used measure of cell-by-cell overlap (Wilson et al. 2005; 2013). The Kappa statistic ranges from −1 (no agreement) to 1 (perfect agreement between layers); a value of 0 indicates random agreement. Overlap is measured using a misclassification matrix that counts how many cells were “wrongly” assigned to each category in both layers, after removing overlap due to chance (Rose et al. 2009). We used the Map Comparison Kit software (Visser & De Nijs 2006) to calculate Kappa; we classified values based on the strength of agreement as low (−1 to 0.2), medium-low (0.2-0.4), medium (0.4-0.6), high (0.6-0.8), and almost perfect (0.8-1) (Ackers et al. 2015).

### d. Trophic level

We estimated changes in time in the mean trophic level 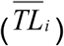 of targeted species for each coastal wetland as recorded in the oral interviews. We estimated the mean trophic level modified from Sala et al. (Sala et al. 2004) for time period *i* by calculating the weighted average of the number of fishers who mentioned a species as a target (*t*) in each site during the participatory mapping, multiplied by the trophic level of the individual species *j*:

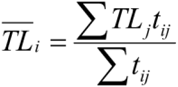

We used an average of the maximum trophic level reported by Spackeen (2009) for Northern Gulf wetlands, based on stable isotope analysis and on the trophic level in FishBase (Froese and Pauly 2014) and SeaLifeBase (Palomares and Pauly 2014) which are based on diet analysis. In some instances, fishers referred to a group, in those cases we used the average trophic level for species in that group.

### d. Perceived species richness and composition

We used participatory drawing, a visual research method to elicit the perceived species richness and composition in coastal wetlands; this method can capture differences in stakeholders’ worldview and bridge linguistic differences in concepts (Gauntlett 2005). Interviewees were presented with a digital diagram depicting a wetland cross section and asked to respond to the question, “what did coastal wetlands look like?” They were then provided with a library of 52 icons depicting common invertebrates, fish, birds, marine mammals, and desert fauna usually found in and around coastal wetlands in the Northern Gulf. Interviewees could then select icons and point to where they should be placed within the wetland diagram. The participatory drawings were created using Adobe Illustrator. Icons came from CEDO’s collection and the image library from the Integration and Application Network, University of Maryland Center for Environmental Science (ian.umces.edu/imagelibrary/).

We used multivariate analysis to describe how patterns of perceived species richness and composition have shifted among time periods and sites. We conducted nonmetric multidimensional scaling (NMDS) based on Bray-Curtis (Sørensen) dissimilarities of the number of mentions for each species in the participatory maps for each site. NMDS provides a graphical representation of perceived species richness and composition by clustering similar site/year combinations. To examine the temporal and site factors, we then constructed biplots using the means of NMDS scores across sites and time periods; we then used multivariate ANOVA with NMDS axis scores as response variables to quantify the spatial and temporal effects.

### e. Environmental history of coastal wetlands

We carried out a bibliographic search for published and unpublished scientific studies that mention coastal wetlands in the Northern Gulf to develop a detailed human-environmental timeline of coastal wetlands in this region. We then generated bibliographic and image databases. Images of coastal wetlands serve as a record of environmental conditions, and include photographs, aerial, and satellite images.

## Results

### Oral interviews

We interviewed 48 resource users; respondent demographic summary characteristics are in Table 3S. Fifty-two percent of interviewees resided in Puerto Penasco, 35% in Ejido Rodolfo Campodonico, and 12% in Golfo de Santa Clara. Sixty-two percent of respondents identified as fishers and 37% as oyster farmers, although 77% of them had also participated in either small-scale or subsistence fisheries during their life. On average, interviewees had 32.1±1.9 years of experience fishing or oyster farming; 67% of respondents were still actively fishing or farming oysters and 33% were retired. Of fishers, 88% of respondents had worked or were still working in small-scale fisheries and 50% in the industrial fleet; only one interviewee mentioned occasional work in sport fishing. Eighty-nine percent of participants confirmed having fished within coastal wetlands. Sixty-four percent of respondents started fishing in the 1970’s or before and 36% in or after the 1980s. Only one participant started fishing in the 2000’s decade. Due to that low sample size, we only consider the decades between 1950-1990 for subsequent analysis.

### Species abundance

Overall, species abundance, suggested by resource user’ responses, showed a consistent downward trend. Figure S2 summarizes uncorrected relative abundance results for the 17 species, as percent of fishers reporting species abundance as high, medium or low in each decade. In all of the species groups investigated, the abundance score was highest in the 1950’s and generally decreased over time. This pattern occurs for most species except spotted sand bass which had lowest reported abundance in the 1970’s and California butterfly ray, which had its highest reported abundance in the 1960’s. Figure 4 shows the output of the fuzzy expert system, where species’ relative abundance was combined with changes in breeding areas, the wetland habitat index (Table 2), and extreme reductions in population size; for most species, there is a consistent downward trend in abundance. We found age-related differences in the abundance trends, the decrease is more pronounced as perceived by resource users that started fishing in the 1950s and 1960s compared to those that began fishing in the 1980s and 1990s.

**Figure 4.**
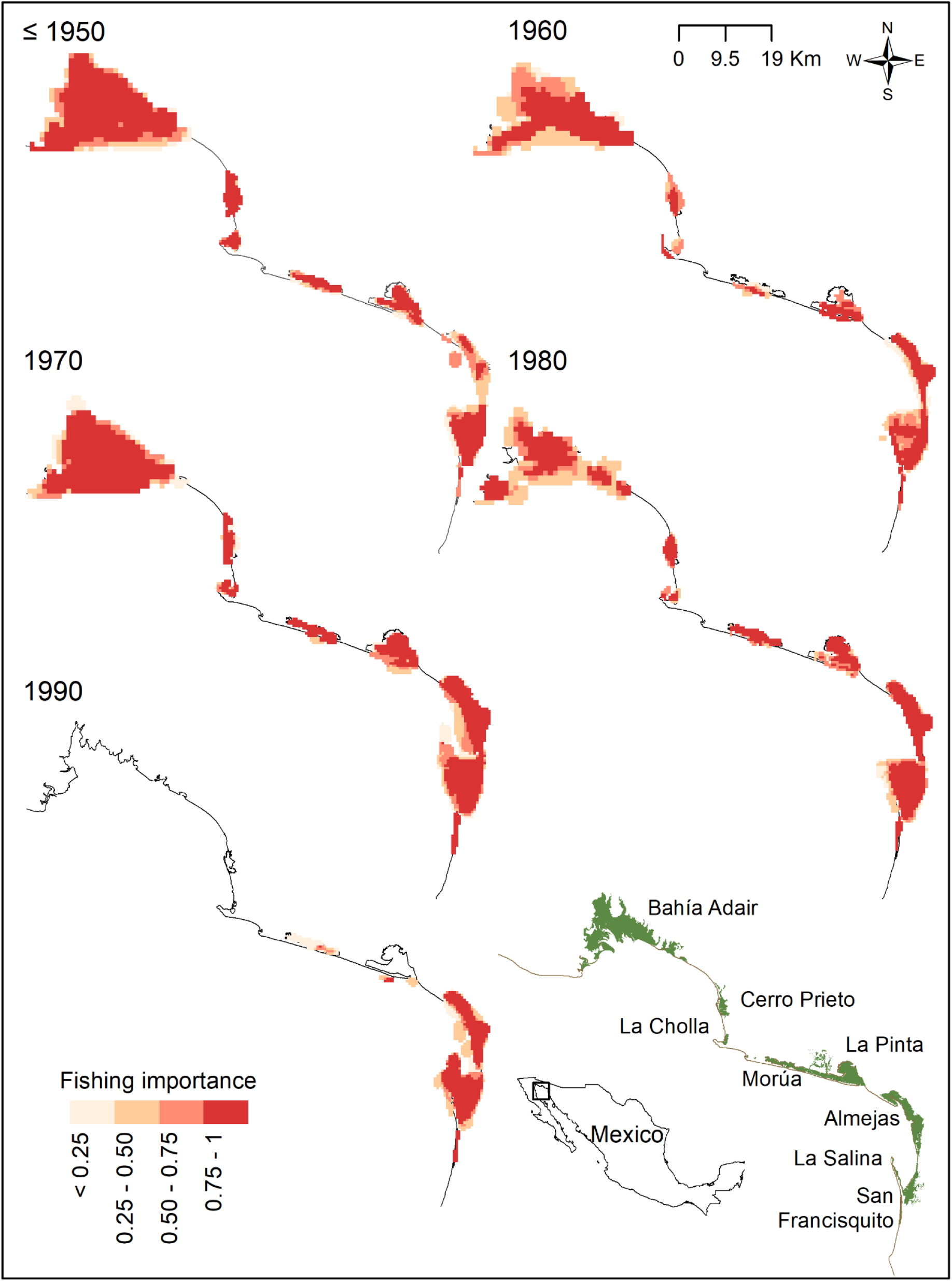
Relative fishing importance index 1950-1990 in coastal wetlands across the coastal corridor in Sonora, Mexico, based on participatory mapping. The index of relative importance was estimated by using a fuzzy sum operator to combine the fuzzy membership function values by fisher and decade for each wetland. For interpretation of the references to color in this figure legend, the reader is referred to the online version of the article.

**Table 2.** Wetland Habitat Index for 1950-2000 and data used to estimate it. Includes the total wetland area converted to other uses, proportion of wetlands lost, and the average proportion of expert users that indicated a loss of wetland functions. Total wetland area in the corridor 46, 300 ha (Glenn et al. 2006).

| Variable | Time period |  |  |  |  |  |
| --- | --- | --- | --- | --- | --- | --- |
|  | 1950 | 1960 | 1970 | 1980 | 1990 | 2000 |
| Total converted (ha) | 45.72 | 45.72 | 65.84 | 99.39 | 399.39 | 799.39 |
| Proportion converted<br>(total wetland area) (PC) | 0.0 | 0.0 | 0.0 | 0.0 | 0.01 | 0.02 |
| Proportion wetlands lost<br>(PL) | 0 | 0 | 0.111 | 0 | 0.075 | 0.026 |
| Average proportion<br>decrease wetland<br>function (DF) | 0 | 0.46 | 0.07 | 0.19 | 0.45 | 0.63 |
| Wetland Habitat Index<br>((PC*0.5)+PL+DF)/3 | 0.00 | 0.15 | 0.06 | 0.06 | 0.18 | 0.22 |

### Areas of fishing importance

Figure 5 shows maps for each wetland for the five decades compared, the areas where the index ~1 are those with the highest fishing importance based on the combined opinion of fishers interviewed. Resource users that started fishing in the 1980s and 1990s indicated fewer areas of fishing importance in a smaller number of wetlands relative to previous decades. Overall, resource users that started fishing in recent decades indicated fewer areas of fishing importance; we found a net decrease when comparing each decade with the subsequent one (Figure 6). The Kappa statistic for assessing agreement between data layers, based on the decade when resource users started fishing, indicated medium to medium-low agreement across decades, illustrating the high variability in areas of fishing importance (Table 3).

**Figure 5.**
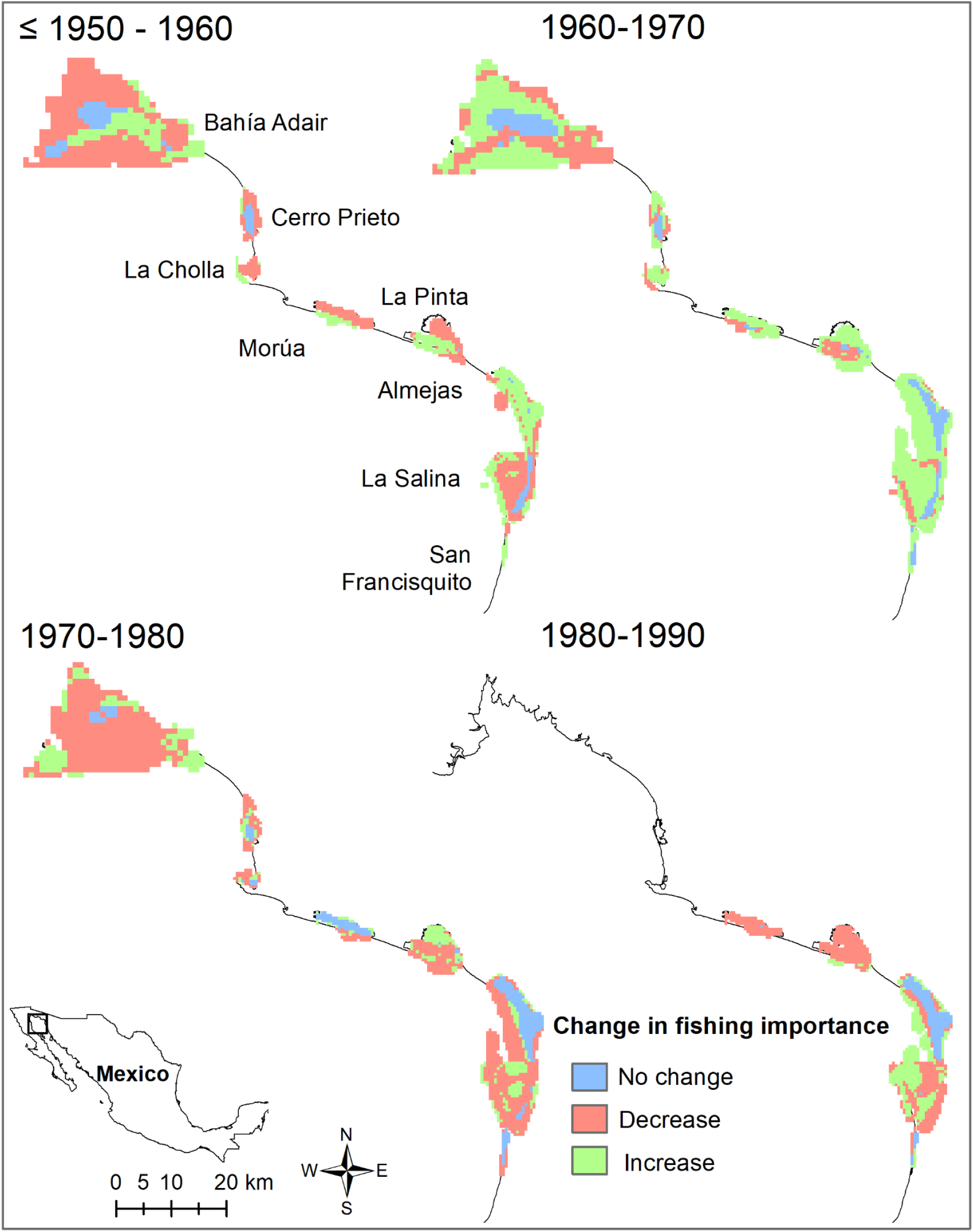
Change in the relative fishing importance index 1950-1990 in coastal wetlands across the coastal corridor in Sonora, Mexico, based on participatory mapping. For interpretation of the references to color in this figure legend, the reader is referred to the online version of the article.

**Figure 6.**
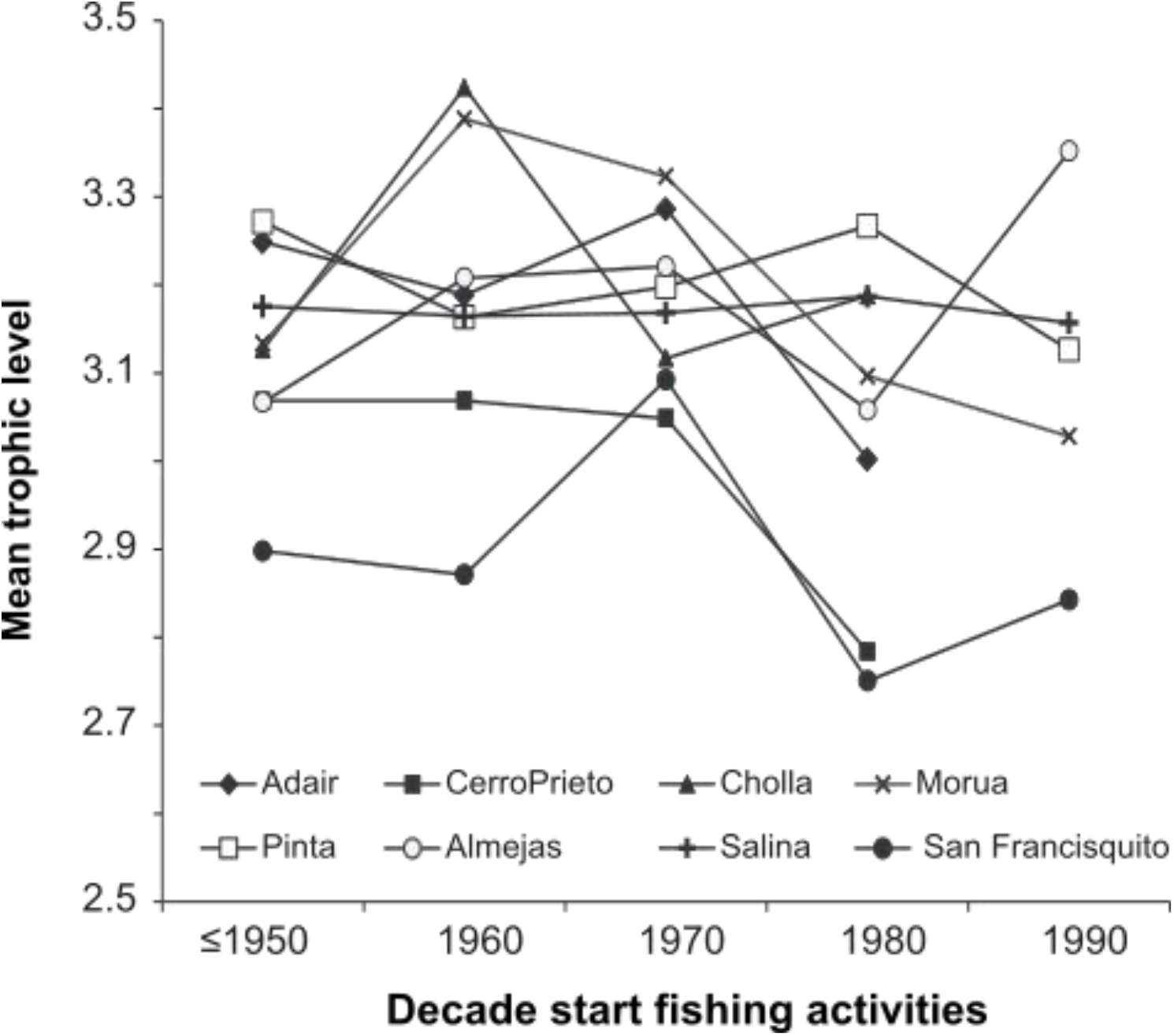
Temporal changes in the mean trophic level of catch for eight coastal wetlands in the Northern Gulf of California based on oral interview data.

**Table 3.** Kappa statistic of overlap between decades when resource users reported fishing for the first time for each coastal wetland. Kappa values express strength of agreement as low (−1 to 0.2), medium-low (0.2-0.4; in **bold**), and medium (0.4-0.6; in italics) (Ackers et al. 2015).

| Wetland | Decade start fishing activities |  |  |  |
| --- | --- | --- | --- | --- |
|  | 50-60 | 60-70 | 70-80 | 80-90 |
| Bahía Adair | *0.425* | *0.492* | **0.310** | - |
| Cerro Prieto | **0.323** | **0.329** | **0.306** | - |
| La Cholla | 0.04 | *0.448* | *0.401* | - |
| Estero Morua |  |  |  |  |
| La Pinta | **0.292** | **0.286** | *0.414* | -0.008 |
| Almejas | **0.297** | **0.309** | **0.398** | **0.486** |
| La Salina | *0.408* | **0.274** | *0.478* | *0.442* |
| San Francisquito | **0.259** | **0.368** | *0.483* | *0.426* |

### Trophic level

We analyzed the mean trophic level of catch for each coastal wetland as recorded in the oral interviews over the five decades studied. The mean trophic level of catch ranged between 3.42 and 2.75 for all sites. The mean trophic level in the 1950’s varied from 3.27 in Bahia Adair to 2.9 in San Francisquito (Figure 7). In the 1960’s six sites experienced either increases in the mean trophic level or small decreases (< 0.05); values then decreased in the 1970’s and 1980’s, but with no clear overall pattern. The decreases between 1970 and 1980, and between 1980 and 1990 are in some cases > 0.15.

**Figure 7.**
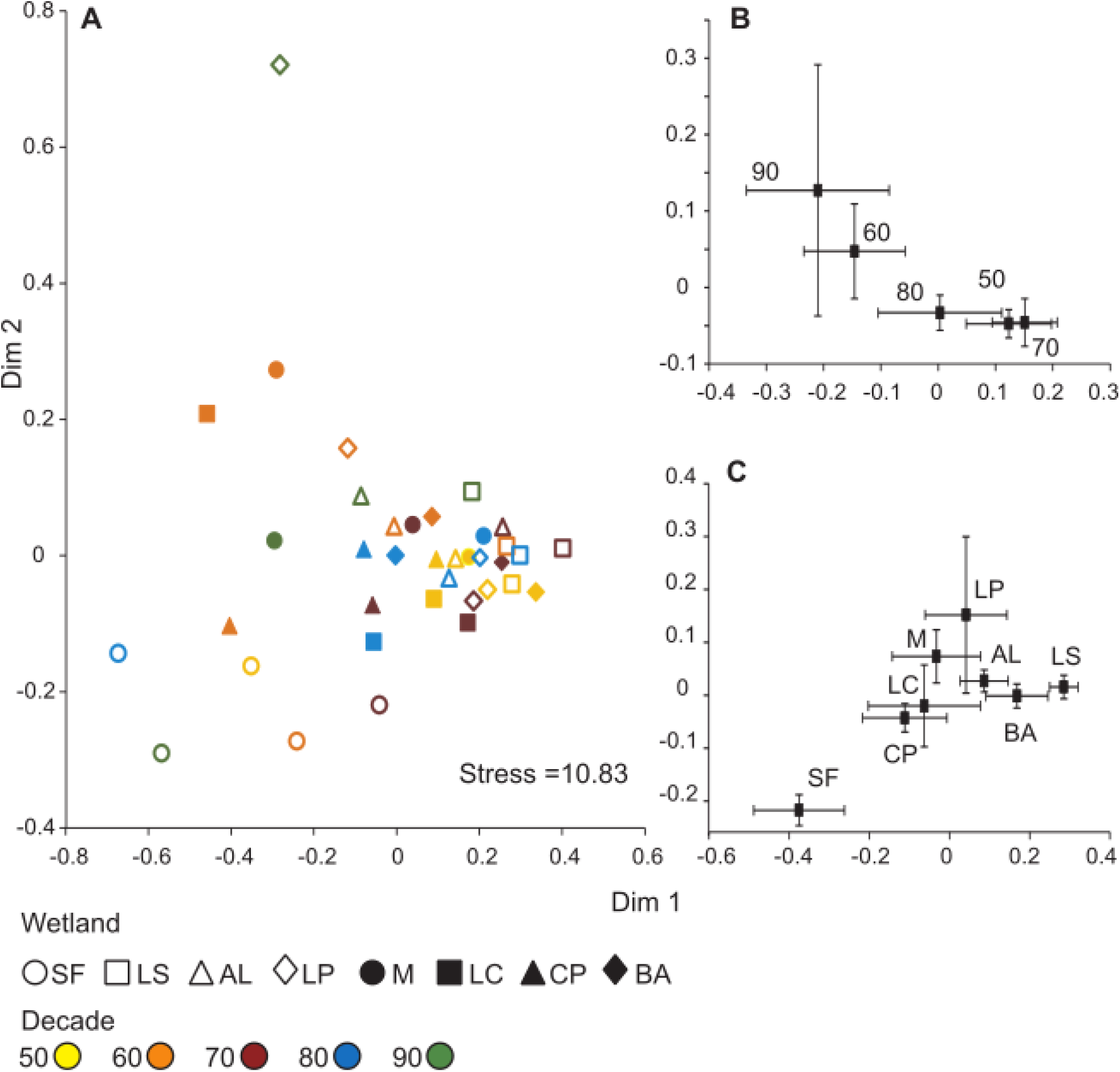
Non-metric multidimensional scaling (A) for perceived species richness and composition data for all sites and time periods. Biplots of non-metric multidimensional scaling showing variation between time periods (B) and sites (C); values are the mean and standard errors for all samples.

### Perceived species richness and composition

Forty interviewees completed the participatory drawing; two participants were partially blind and two stated they did not know the systems sufficiently to complete the drawing. In five cases, drawings were not completed because we either experienced technical difficulties or ran out of time during the interview. Participatory drawings (i.e. Figure S3; full set available from the data repository (https://knb.ecoinformatics.org/view/doi%3A10.5063%2FF12R3Q48) illustrate the different habitats within coastal wetlands and biodiversity. All fifty-two taxa were portrayed in the drawings. Time since the participant started fishing was negatively correlated with the number of species placed in a drawing, likely due to increasing familiarity with technology in younger fishers.

The ordination of perceived species richness and composition (Figure 8) revealed that there was no clear pattern between decades or sites; in Figure 8A, tightly clustered sites have similar perceived richness and composition. La Salina was the most tightly clustered site, where perceived richness and composition varied the least across decades. The wetlands of San Francisquito and La Salina had the least overlap in perceived richness and composition with other wetlands. Site was the only significant variable for the ANOVA on the scores of the NMDS axis 1 and 2 (Table 4). There was no effect of year (Figure 8B) and no significant interaction between year and site. The biplot of NMDS scores (Figure 8C) for sites shows a clear gradient in perceived species richness and composition relative to wetland size from the two largest sites (La Salina and Bahia Adair), to one of the smallest sites, San Francisquito, which is also the most tidally-restricted.

**Figure 8.**
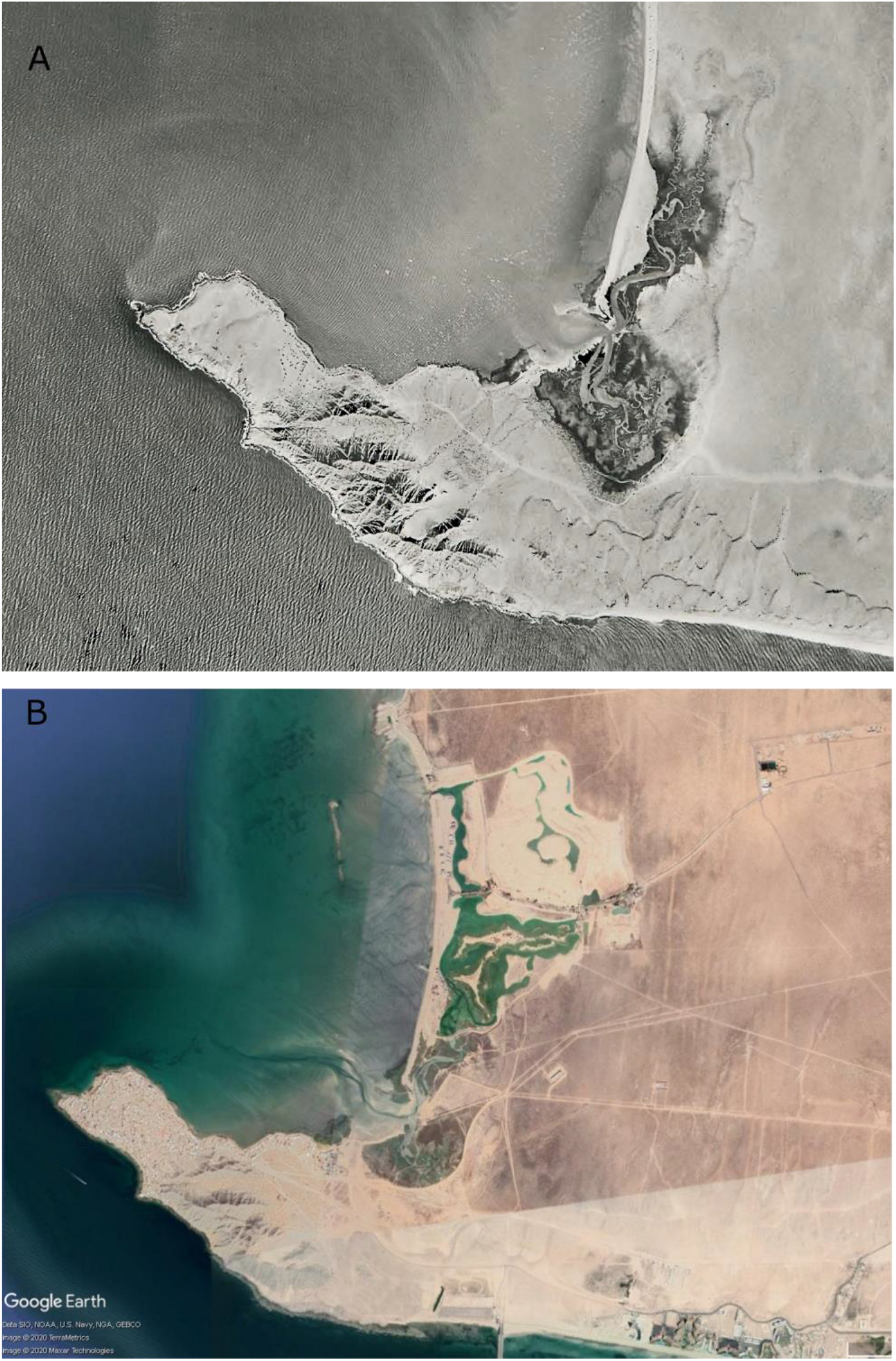
Estero La Cholla, Puerto Peñasco Coastal Corridor in Sonora, México. A. 1956. Joseph F. Schreiber Jr. B. Image composite from Google Earth. Data SIO, NOAA, U.S. Navy, NGA, GEBCO, Image © TerraMetrics. Image @ 2020 Maxar technologies. Retrieved January 2020.

**Table 4.**
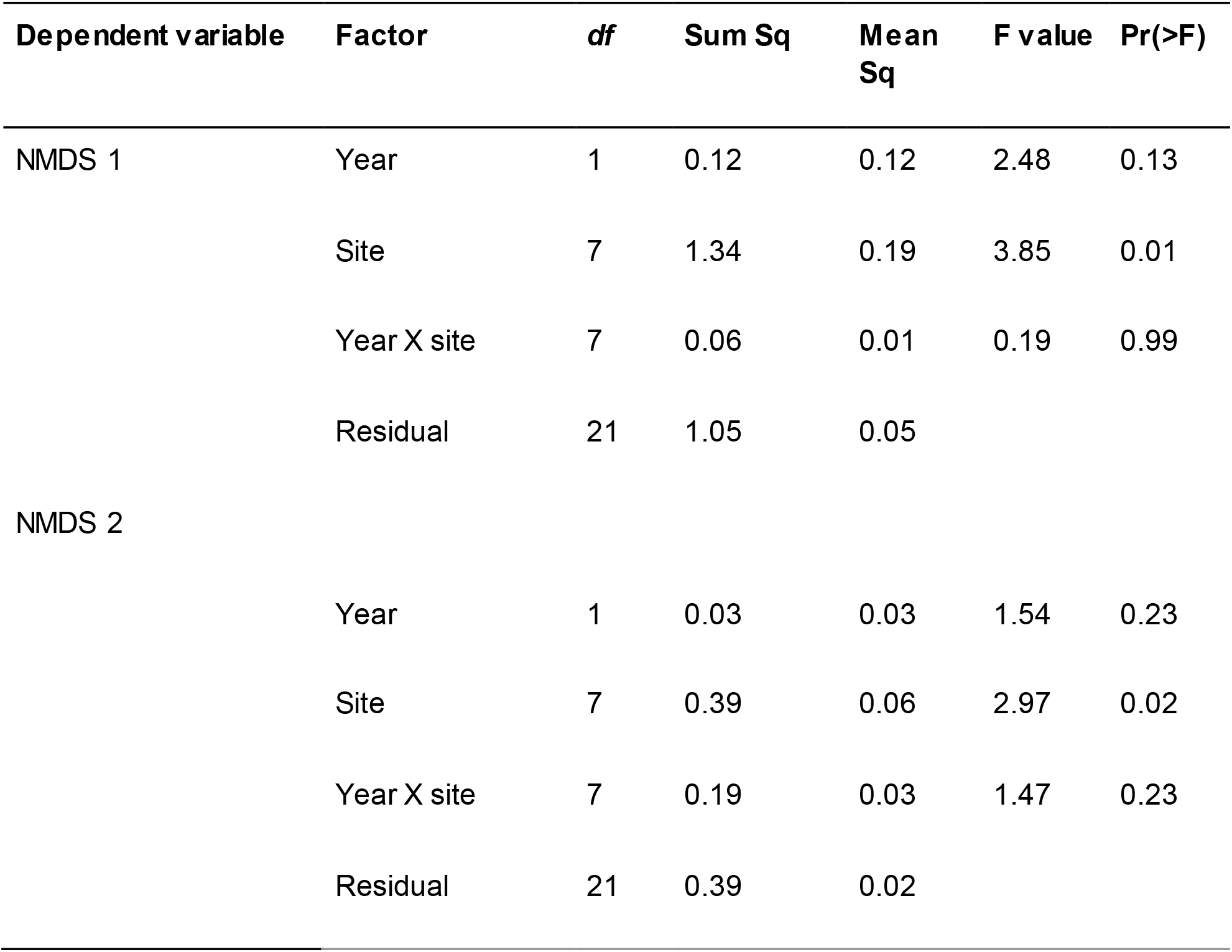
Results of ANOVAs quantifying the effects of year and site on the species composition non metric multidimensional scaling scores.

### Environmental history of coastal wetlands

We identified 140 bibliographic references with information on wetlands in the Northern Gulf, including articles describing scientific research. These references are provided in Table 2S, and serve as a record of existing ecological knowledge in these sites. The references are also available as a BibTeX file from the data repository (https://knb.ecoinformatics.org/view/doi%3A10.5063%2FF12R3Q48). We used this information to describe the environmental history of coastal wetlands in the Northern Gulf of California; we developed a detailed timeline from 5000 BPE to 2020 with key environmental factors, landscape impacts, social and environmental drivers of change, and the resulting implications for wetland function and structure (Figure S4). A detailed environmental history is in Text S6. Briefly, the current configuration of coastal wetlands formed during the Holocene (5000 BP), when the sea level rise reached its highest point (Ortlieb 1991). These wetlands were used for thousands of years by humans; archeological evidence confirms human use as far back as the Middle Archaic (5500 – 3500 BP) (Foster et al. 2008). Anthropogenic impacts increased in the 20th century following modern occupation and increased fishing activities (Munro Palacio 2007). Tourism development has had the most profound impact on coastal wetlands in this region, as it has involved filling wetland areas and conversion to other uses (Glenn et al. 2006). Currently, land tenure surrounding the wetlands in the Puerto Peñasco coastal corridor includes communal and private property and land use includes tourism, residential, fishing camps, and aquaculture (Morzaria-Luna et al. 2014a). Wetlands in Mexico are public domain and are administered as part of the Maritime-Terrestrial Federal Zone, which extends up to 20 meters inland from the highest tide line (Cortina-Segovia et al. 2007). Coastal wetlands in this region are now under different conservation frameworks, including Ramsar sites (Morzaria-Luna et al. 2014a) but are still threatened by human development.

We also gathered a collection of 100 photographs, aerial and remote sensing images taken between 1965 and 2008. These images provide a baseline of the original conditions, extent and habitats of coastal wetlands in the Northern Gulf. These photographs document extensive changes in coastal wetlands such as La Cholla (Figure 3), where a large area of wetland was converted into tourism development. The aerial photos include those taken by CEDO staff and a collection of photos taken during the 1950 and 60’s and donated to CEDO by Joseph F. Schreiber Jr. (1932-2015), professor emeritus in the Department of Geosciences at the University of Arizona. The images cover the coastline between La Salina (31°29’53’’ N’ 114°09’52” W) to the North and Estero San Francisquito (31°53’50 N; 113°06’09” W) to the south. We deposited the image inventory and associated images in the data repository (https://knb.ecoinformatics.org/view/doi%3A10.5063%2FF12R3Q48); images subject to copyright could not be included in the repository.

## Discussion

We documented historical patterns of commercial species abundance, areas of fishing importance, trophic level, and species richness and composition in coastal wetlands in the Northern Gulf of California, Mexico, across sites and time periods. Our main sources of information was the Local Ecological Knowledge (LEK) of wetland resource users (small-scale fishers and oyster farmers) and expert interviews. As we show here, Local ecological knowledge (LEK) can be used to inform how the ecosystem might have been structured decades ago, before time-series data began for most species, and can supplement scientific, archeological, and naturalist records, particularly when dealing with data-limited species (Ainsworth and Pitcher 2005). We elicited LEK from participants through oral histories, participatory mapping, and participatory illustration. We also reconstructed the environmental history of coastal wetlands in this region from bibliographic sources and photographic records, to document impacts that could have affected coastal fisheries. LEK, in conjunction with historical data from diverse information sources can help set conservation and management goals (McClenachan et al. 2012). In this case, documenting small-scale fisheries’ use of coastal wetlands in the Northern Gulf of California can offer an important perspective for the management of natural resources in the region (Lozano-Montes et al. 2008) and can help assess the full implications of the loss of ecosystem services from overfishing (Zu Ermgassen et al. 2012). Our study had a large enough sample size relative to what is considered necessary to make robust conclusions derived from fishers’ memories (≥30 interviewees) (Sáenz-Arroyo and Revollo-Fernández 2016).

### Trends in target species’ relative abundance indicate shifting baselines

We found that for most species abundance decreased, based on resource user responses, and there were age-related differences in the abundance trends, with a more pronounced decrease for resource users that started fishing in the 1950s and 1960s compared to later years. This suggests that younger resource users have not experienced the biodiversity and fish abundance of the Gulf of California that their parents experienced, so this knowledge is being lost from collective memory (Bunce et al. 2008). Shifting baselines describes the phenomenon where historical environmental information is lost over time and ecosystem changes go unrecognized because each successive generation judges the environmental state against a current reference point (Sáenz-Arroyo et al. 2005a; Jones et al. 2020), resulting on different expectations of a ‘desirable’ environment, increased tolerance for environmental degradation, and inappropriate baselines for management and conservation (Soga and Gaston 2018). Previous research has found evidence of changing perceptions of fish abundance and biodiversity in the Gulf of California. Lozano-Montes et al. (2008) found that older fishers reported a higher loss of fishing sites and higher abundance over the past 50 years relative to middle-aged and young fishers. While Sáenz-Arroyo et al. (2005a) reported that the number of fishing sites and fish species that younger fishers considered depleted was approximately one-quarter of that reported by older fishers.

We confirmed that LEK can be a source of high-quality and low-cost information about species abundance (Anadón et al. 2009). Local Ecological Knowledge has previously been used to obtain qualitative and quantitative estimates of fishery species’ abundance or catch trends (Ainsworth et al. 2008b; Lozano-Montes et al. 2008; Ainsworth 2011; Beaudreau et al. 2011; Sáenz-Arroyo and Revollo-Fernández 2016). LEK appears best suited to detect changes over long time periods; abundance trends based on LEK correlate poorly with observed data in species that exhibit high inter-annual variability (Ainsworth and Pitcher 2005). It is important to note that LEK perceptions of resource status may incorporate subjective opinions regarding how those populations are managed or allocated (Beaudreau and Levin 2013). Our abundance trends (Figure 4) coincide with previous research in the Gulf of California. Ainsworth (2011) found a significant decline in the abundance of fished and unfished species since the 1950’s including turtles, elasmobranchs, penaeid shrimps, Gulf Coney, and totoaba. While, Lozano-Montes et al. (2008) found that fishers perceived that fishery resources declined by over 60% between 1950-2000.

### Important fishing areas decreased over time

We found that the spatial extent of areas of fishing importance, as perceived by resource users, shrunk, with smaller areas in fewer wetlands (Figures 5–6). Small-scale fishers usually operate within a limited range, and use low-technology methods and small and/or traditional fishing vessels and gears (Wanyonyi et al. 2018). The locations where fishers fish are influenced by a wide range of biological, regulatory, economic, and cultural factors including abundance (Gillis et al. 1993), environmental conditions, economic variables, regulatory constraints (Naranjo-Madrigal et al. 2015), such as the presence of other fishers (Daw 2008), travel costs, logistical and technical considerations (Duberstein 2009), depth or distance from shore (Coll et al. 2014), and regulations that limit fishing in certain areas, such as Marine Protected Areas (Beaudreau and Whitney 2016). The reductions in areas of fishing importance could be attributed to these socioeconomic impacts or to perceived reductions in fish abundance as a result of overfishing and environmental degradation.

We expected to find evidence of the effects of overfishing when documenting areas of fishing importance within coastal wetlands. These areas are mainly used by small-scale coastal fisheries and multidecadal declines (Sáenz-Arroyo et al. 2005b) and serial depletions (Luquin-Covarrubias et al. 2022) in coastal fisheries are attributed to overfishing (Sagarin et al. 2008). The imbalance in species interactions-predator-prey relationships caused by overfishing can extend beyond the direct effects on community structure; overfishing can also reduce species richness, diversity, and distribution (Kennett et al. 2008; Gedan et al. 2009) and have a bottom-up effect in the food web to the level of nutrients and system biogeochemistry (Coleman and Williams 2002; Scheffer et al. 2005), preconditioning the system for further impacts from eutrophication, disease outbreaks, and toxic blooms (Jackson et al. 2001).

### Trophic level showed no temporal or spatial pattern

We did not find a decreasing trend in the mean trophic level of catch (Figure 6), as has been previously documented for worldwide fishery landings (Pauly et al. 1998a) and for coastal fisheries in the Southern Gulf of California (Sala et al. 2004). The latter found a decrease in the mean trophic level of catch, from 4.2 to 3.8 between 1970 and 2000. We did however find decreases between 1970 and 1980 that were > 0.15, which can be considered ecologically significant (Essington et al. 2006), as this decrease would represent an ≈50% decline in the primary production required to sustain a given amount of catch (Pauly and Christensen 1995).

In general, decreases in trophic levels can result from overfishing species from higher trophic levels and their substitution with lower-trophic level fisheries (Pauly et al. 1998b), or from the sequential addition of lower trophic level fisheries while fisheries for higher trophic-level species are maintained (Essington et al. 2006). In the case of Northern Gulf fisheries, our results suggest the latter, that new species from both high and low trophic levels were subsequently targeted across time periods, resulting in small changes in trophic level. This result coincides with the historical development of small-scale fisheries in the region, which in the past 50 years has gone from low-impact, targeting few species, to a high-impact activity where a large diversity of fishing gears is used target over 70 species of fish and invertebrates (Lluch-Cota et al. 2007; Cudney-Bueno et al. 2009).

### Impacts on coastal wetlands in the Northern Gulf of California threaten ecosystem function

The degradation and destruction of natural habitats, including coastal wetlands, has increased in recent decades but has been prevalent throughout human history (Zu Ermgassen et al. 2012). This has motivated an expansion in the conservation of remaining natural areas and restoration of damaged habitats to maintain ecosystem services (Zedler and Kercher 2005). We documented dramatic shifts in the spatial extent of areas of fishing importance and in the abundance of target species within coastal wetlands in the Northern Gulf. Habitat change could account for some of the changes in species abundance; we documented in the environmental history of wetlands in this region (Text 1S) that they have been affected by tidal restriction, which limits tidal influence, and habitat degradation and conversion, impacting or eliminating the ecosystem services provided by coastal wetlands, including carbon and nutrient sequestration, coastal defense, and wildlife habitat (Gedan et al. 2009). Although different conservation mechanisms have been applied to protect wetlands in the Northern Gulf, challenges remain such as insufficient enforcement of existing environmental legislation and low economic penalties for wetland modification or destruction (Morzaria-Luna et al. 2014a). It is also likely that increasing fishing pressure in marine areas outside coastal wetlands has affected wetland assemblage structure; predator removal from overfishing can trigger trophic cascades that may even affect wetland vegetation (Altieri et al. 2012).

We found a gradient in perceived species richness and composition relative to wetland size. This species-area relationship has been shown to be a general pattern found in ecology; and can be explained by the fact that a large area can contain more habitats, species, and individuals of each species (Gray 2001).

## Our results reflect the transformation of fishing communities

The richness of marine resources in the Northern Gulf of California and in the Puerto Peñasco corridor is reflected in its long history of fisheries exploitation. Coastal communities went from isolated groups that traded sea shells with indigenous communities up north to well established coastal towns focused on high-value fisheries (Mitchell and Foster 2000; Rodríguez-Quiroz et al. 2010; Mitchell et al. 2015). Local fisheries transformed from using basic equipment to more technologically advanced gear. The increased fishing effort and modification of wetlands led to a decrease in species abundances within the more scarce and smaller fishing grounds. These changes have reflected in local communities where fisheries are perceived as an important part of the local identity (Enríquez Acosta et al. 2015). The interest of local fishers in the state of fisheries have led to self-organization processes (McLain et al. 2018). Local communities from the Puerto Peñasco corridor have collaborated with NGOs and research groups to improve the management of their fisheries resources (Turk-Boyer et al. 2014a; Munguía-Vega et al. 2015). Within this collaborative management strategies historical information is key to determine wetlands baselines and set recovery goals.

## Conclusions

We have created a permanent record of the perceptions of resource users that exploited fishery resources in the Northern Gulf following the modern establishment of fishing towns. This is significant because fishers’ LEK is getting eroded as fisheries livelihoods change, due to migration, education, transition to other occupations, diminished capacity for sustainable fishing, loss of access to traditional resources, and vulnerability to the COVID-19 pandemic (Kai et al. 2014; Lopez-Ercilla et al. 2021).

Our analysis of historical patterns of commercial species abundance, areas of fishing importance, trophic level, and species richness and composition in coastal wetlands in the Northern Gulf of California, Mexico, can be used to estimate historical ecosystem services and to set targets for ecological restoration, management practices and conservation efforts, which depend on a knowledge of past abundance (Lozano-Montes et al. 2008; Zu Ermgassen et al. 2012). Baseline ecological studies in the Northern Gulf do not exist prior to the 1970’s (Lozano-Montes et al. 2008) and human impacts on coastal wetlands in this region were not documented until the 1990’s (Valdes-Casillas et al. 1999). Documenting past abundance trends and fishing areas is also significant because the shifting environmental baselines introduced by climate change will create new challenges when trying to measure ecological thresholds or establish management goals in coastal wetlands (Morzaria-Luna et al. 2014b; Hirsch 2020), because the rate of climate change is faster, nonlinear, can generate feedback and forward loops and thresholds of irreversible change, and will likely lead to a novel future (Fernández-Llamazares et al. 2015).

## Author contributions statement

H. Morzaria-Luna conceptualized the project and secured funding. M. Urquidi and Sánchez Cruz carried out interviews. M. Urquidi, H. Morzaria-Luna, J.M. Dorantes Hernández processed data. H. Morzaria-Luna, M. Urquidi, G. Cruz-Piñon, and J.M. Dorantes carried out the data analysis. H. Morzaria-Luna, J.M. Dorantes, G. Cruz-Piñon, and P. Valdivia Jiménez generated figures and tables. H. Morzaria-Luna and J.M. Dorantes generated the initial manuscript. All authors contributed to the final mansucript and reviewed the final version.

## Data availability statement

The datasets generated are available in the KNB repository https://knb.ecoinformatics.org/view/doi%3A10.5063%2FF12R3Q48, personally identifiable information for interviewees has been removed and data have been aggregated as necessary.

## Funding

This project was funded by a Mia J. Tegner Memorial Research Grant in Marine Environmental History and Historical Marine Ecology from the Marine Conservation Biology Institute. Additional funding was provided by The David and Lucile Packard Foundation, grant 2006-30328 and The Sandler Family Supporting Foundation to CEDO Intercultural.

## Conflict of Interest

The authors declare that they have no conflict of interest.

## Acknowledgments

This project was funded by a Mia J. Tegner Memorial Research Grant in Marine Environmental History and Historical Marine Ecology from the Marine Conservation Biology Institute. Additional funding was provided by the The David and Lucile Packard Foundation, grant 2006-30328 and The Sandler Family Supporting Foundation to CEDO Intercultural. C. Alva, E. López, and A. Iris Maldonado assisted with the interviews and participatory mapping. R. Loaiza and J.R. Sanchez provided information on fishing areas. C. Ainsworth provided guidance on the application of the fuzzy logic approach. K. Flessa and P. Nagler provided access to maps and images of coastal wetlands.

## Notes

https://knb.ecoinformatics.org/view/doi%3A10.5063%2FF12R3Q48

### Competing Interest Statement

The authors have declared no competing interest.

## References

Ackers SH, Davis RJ, Olsen KA, Dugger KM (2015) The evolution of mapping habitat for northern spotted owls (Strix occidentalis caurina): A comparison of photo-interpreted, Landsat-based, and lidar-based habitat maps. Remote Sens Environ 156:361–373

Ainsworth CH (2011) Quantifying species abundance trends in the Northern Gulf of California using Local Ecological Knowledge. Mar Coast Fish 3:190 – 218

Ainsworth CH, Pitcher TJ (2005) Using local ecological knowledge in ecosystem models. Fisheries assessment and management in data-limited situations Alaska Sea Grant College Program, University of Alaska Fairbanks, Fairbanks, Alaska, USA 289–304

Ainsworth CH, Pitcher TJ, Heymans JJ, Vasconcellos M (2008a) Reconstructing historical marine ecosystems using food web models: Northern British Columbia from Pre-European contact to present. Ecol Modell 216:354–368

Ainsworth CH, Pitcher TJ, Rotinsulu C (2008b) Evidence of fishery depletions and shifting cognitive baselines in Eastern Indonesia. Biol Conserv 141:848–859

Altieri AH, Bertness MD, Coverdale TC, et al (2012) A trophic cascade triggers collapse of a salt-marsh ecosystem with intensive recreational fishing. Ecology 93:1402–1410

Álvarez-Romero JG, Pressey RL, Ban NC, et al (2013) Marine conservation planning in practice: lessons learned from the Gulf of California. Aquat Conserv. https://doi.org/10.1002/aqc.2334

Alvirde-López SB (2012) Estimación de la red trófica en el Estero Morúa, Sonora, México, un humedal costero del norte del Golfo de California

Ames T (2007) Putting fishers’ knowledge to work: Reconstructing the Gulf of Maine cos spawning grounds on the basis of local ecological knowledge. Fishers’ Knowledge in Fisheries Science and Management 353–363

Anadón JD, Giménez A, Ballestar R, Pérez I (2009) Evaluation of local ecological knowledge as a method for collecting extensive data on animal abundance. Conserv Biol 23:617–625

Arce-Acosta M, Ramírez-Rodríguez M, De-la-Cruz-Agüero G (2018) Small scale fisheries operative units in the west central region of the Gulf of California, Mexico. Ocean Coast Manag 160:58–63

Azzurro E, Moschella P, Maynou F (2011) Tracking signals of change in Mediterranean fish diversity based on local ecological knowledge. PLoS One 6:e24885

Basurto X, Bennett A, Weaver AH, et al (2013) Cooperative and noncooperative strategies for small-scale fisheries’ self-governance in the globalization era: implications for conservation. Ecol Soc 18:

Baum JK, Myers RA (2004) Shifting baselines and the decline of pelagic sharks in the Gulf of Mexico. Ecol Lett 7:135–145

Beaudreau AH, Levin PS (2013) Advancing the use of local ecological knowledge for assessing data-poor species in coastal ecosystems. Ecol Appl. https://doi.org/10.1890/13-0817.1

Beaudreau AH, Levin PS, Norman KC (2011) Using folk taxonomies to understand stakeholder perceptions for species conservation. Conservation Letters 4:451–463

Beaudreau AH, Whitney EJ (2016) Historical patterns and drivers of spatial changes in recreational fishing activity in Puget Sound, Washington. PLoS One 11:e0152190

Brusca RC, Cudney-Bueno R, Moreno-Báez M (2006) Gulf of California esteros and estuaries.Analysis, state of knowledge, and conservation and priority recommendations. Arizona-Sonora Desert Museum

Bunce M, Rodwell LD, Gibb R, Mee L (2008) Shifting baselines in fishers’ perceptions of island reef fishery degradation. Ocean Coast Manag 51:285–302

Calderon-Aguilera LE, Marinonea SG, Aragon-Noriega EA (2003) Influence of oceanographic processes on the early life stages of the blue shrimp (*Litopenaeus stylirostris*) in the Upper Gulf of California. J Mar Syst 39:117–128

Cánovas-Molina A, García-Charton JA, García-Frapolli E (2021) Assessing the contribution to overfishing of small-and large-scale fisheries in two marine regions as determined by the weight of evidence approach. Ocean Coast Manag 213:105911

Chan MN, Beaudreau AH, Loring PA (2019) Exploring diversity in expert knowledge: variation in local ecological knowledge of Alaskan recreational and subsistence fishers. ICES J Mar Sci 76:913–924

Cisneros-Mata MA (2010) The importance of fisheries in the Gulf of California and ecosystem-based sustainable co-management for conservation. In: Brusca R (ed) Gulf of California:Biodiversity and Conservation. University of Arizona Press; Arizona-Sonora Desert Museum, Tucson Ariz., pp 119–134

Cisneros-Montemayor AM, Vincent ACJ (2016) Science, society, and flagship species: social and political history as keys to conservation outcomes in the Gulf of California. Ecol Soc 21:art9

Close CH, Brent Hall G (2006) A GIS-based protocol for the collection and use of local knowledge in fisheries management planning. J Environ Manage 78:341–352

Cohen J (1960) A coefficient of agreement for nominal scales. Educ Psychol Meas 20:37–46

Coleman FC, Williams SL (2002) Overexploiting marine ecosystem engineers: potential consequences for biodiversity. Trends Ecol Evol 17:40–44

Coll M, Carreras M, Ciércoles C, et al (2014) Assessing fishing and marine biodiversity changes using fishers’ perceptions: the Spanish Mediterranean and Gulf of Cadiz case study. PLoS One 9:e85670

Coll M, Lotze HK (2016) Ecological Indicators and Food-Web Models as Tools to Study Historical Changes in Marine Ecosystems. In: Schwerdtner Máñez K, Poulsen B (eds)Perspectives on Oceans Past. Springer Netherlands, Dordrecht, pp 103–132

Cortina-Segovia S, Brachet-Barro G, Ibañez de la Calle M, Quiñones-Valades L (2007) Océanos y costas. Análisis del marco jurídico e instrumentos de política ambiental en México. Secretaría de Medio Ambiente y Recursos Naturales. Instituto Nacional de Ecología, México, D.F.

Cudney-Bueno R, Bourillón L, Sáenz-Arroyo A, et al (2009) Governance and effects of marine reserves in the Gulf of California, Mexico. Ocean Coast Manag 52:207–218

Cudney-Bueno R, Turk Boyer PJ (1998) Pescando entre mareas del Alto Golfo de California:Una guía sobre pesca artesanal, su gente y sus propuestas de manejo. CEDO Intercultural, A.C., Puerto Peñasco, Sonora. México

Daw TM (2008) Spatial distribution of effort by artisanal fishers: Exploring economic factors affecting the lobster fisheries of the Corn Islands, Nicaragua. Fish Res 90:17–25

DeCelles GR, Martins D, Zemeckis DR, Cadrin SX (2017) Using Fishermen’s Ecological Knowledge to map Atlantic cod spawning grounds on Georges Bank. ICES J Mar Sci 74:1587–1601

DOF (2013) NORMA Oficial Mexicana NOM-002-SAG/PESC-2013, Para ordenar el aprovechamiento de las especies de camarón en aguas de jurisdicción federal de los Estados Unidos Mexicanos. Diario Oficial de la Federación. 11/07/2013

Duberstein JN (2009) The shape of the commons: Social networks and the conservation of small-scale fisheries in the Northern Gulf of California, Mexico. Ph.D., School of Natural Resources and Environment

Enríquez-Acosta JÁE (2008) Las nuevas ciudades para el turismo. Caso Puerto Peñasco, Sonora, México. Barcelona. Universidad de Barcelona

Enríquez Acosta JÁ, Mayorquín HH, Sarabia CL (2015) Percepciones de los habitantes acerca de la actividad turística, la crisis económica y los problemas sociales en Puerto Peñasco, México. TURyDES 8:

Essington TE, Beaudreau AH, Wiedenmann J (2006) Fishing through marine food webs. Proc Natl Acad Sci U S A 103:3171–3175

Fernández-Llamazares Á, Díaz-Reviriego I, Luz AC, et al (2015) Rapid ecosystem change challenges the adaptive capacity of Local Environmental Knowledge. Glob Environ Change 31:272–284

Foster MS, Mitchell DR, Huckleberry G, Dettman DL (2008) Observations on the archaeology, paleoenvironment, and geomorphology of the Puerto Peñasco area of northern Sonora, Mexico. Kiva 73:263–290

Frans VF, Augé AA (2016) Use of local ecological knowledge to investigate endangered baleen whale recovery in the Falkland Islands. Biol Conserv 202:127–137

Friedman M, Kandel A (1999) Introduction to pattern recognition. World Scientific Publishing Company

Froese R, Pauly D (eds) (2014) FishBase. World Wide Web electronic publication. www.fishbase.org, (11/2014)

Gauntlett D (2005) Using creative visual research methods to understand media audiences. MedienPädagogik: Zeitschrift für Theorie und Praxis der Medienbildung 9:1–32

Gedan KB, Silliman BR, Bertness MD (2009) Centuries of human-driven change in salt marsh ecosystems. Ann Rev Mar Sci 1:117–141

Gillis DM, Peterman RM, Tyler AV (1993) Movement Dynamics in a Fishery: Application of the Ideal Free Distribution to Spatial Allocation of Effort. Can J Fish Aquat Sci 50:323–333

Glenn EP, Nagler PL, Brusca RC, Hinojosa-Huerta O (2006) Coastal wetlands of the northern Gulf of California: inventory and conservation status. Aquatic Conservation-Marine and Freshwater Ecosystems 16:5–28

Gomez-Sapiens MM, Soto-Montoya E, Hinojosa-Huerta O (2013) Shorebird abundance and species diversity in natural intertidal and non-tidal anthropogenic wetlands of the Colorado River Delta, Mexico. Ecol Eng 59:74–83

Goodman LA (1961) Snowball Sampling. Ann Math Stat 32:148–170

Gray JS (2001) Marine diversity: the paradigms in patterns of species richness examined. scimar 65:41–56

Gutiérrez NL, Hilborn R, Defeo O (2011) Leadership, social capital and incentives promote successful fisheries. Nature 470:386–389

Hall GB, Close CH (2007) Local knowledge assessment for a small-scale fishery using geographic information systems. Fish Res 83:11–22

Hirsch SL (2020) Anticipatory practices: Shifting baselines and environmental imaginaries of ecological restoration in the Columbia River Basin. Environment and Planning E: Nature and Space 3:40–57

Jackson JB, Kirby MX, Berger WH, et al (2001) Historical overfishing and the recent collapse of coastal ecosystems. Science 293:629–637

Jones BL, Unsworth RKF, Udagedara S, Cullen-Unsworth LC (2018) Conservation Concerns of Small-Scale Fisheries: By-Catch Impacts of a Shrimp and Finfish Fishery in a Sri Lankan Lagoon. Frontiers in Marine Science 5.: https://doi.org/10.3389/fmars.2018.00052

Jones LP, Turvey ST, Massimino D, Papworth SK (2020) Investigating the implications of shifting baseline syndrome on conservation. People and Nature 2:1131–1144

Kai Z, Woan TS, Jie L, et al (2014) Shifting baselines on a tropical forest frontier: extirpations drive declines in local ecological knowledge. PLoS One 9:e86598

Kennett DJ, Voorhies B, Wake TA, Martinez N (2008) Long-term effects of human predation on marine ecosystems in Guerrero, Mexico. In: Rick TC, Erlandson JM (eds) Human Impacts on Ancient Marine Ecosystems: A Global Perspective. University of California Press, Berkeley and Los Angeles, California, pp 103–124

Lanz E, Nevárez-Martínez MO, López-Martínez J, Dworak JA (2008) Spatial distribution and species composition of small pelagic fishes in the Gulf of California. Rev Biol Trop 56:575–590

Leduc AOHC, De Carvalho FHD, Hussey NE, et al (2021) Local ecological knowledge to assist conservation status assessments in data poor contexts: a case study with the threatened sharks of the Brazilian Northeast. Biodivers Conserv 30:819–845

Lluch-Cota SE, Aragón-Noriega EA, Arreguín-Sánchez F, et al (2007) The Gulf of California:review of ecosystem status and sustainability challenges. Prog Oceanogr 73:1–26

Lopez-Ercilla I, Espinosa-Romero MJ, Fernandez Rivera-Melo FJ, et al (2021) The voice of Mexican small-scale fishers in times of COVID-19: Impacts, responses, and digital divide. Mar Policy 131:104606

Lotze HK (2010) Historical Reconstruction of Human-Induced Changes in U.S. Estuaries. Oceanography and Marine Biology 267–338

Lotze HK, Lenihan HS, Bourque BJ, et al (2006) Depletion, Degradation, and Recovery Potential of Estuaries and Coastal Seas. Science 312:1806–1809

Lozano-Montes H, Pitcher TJ, Haggan N (2008) Shifting environmental and cognitive baselines in the upper Gulf of California. Front Ecol Environ 6:75–80

Luquin-Covarrubias MA, Morales-Bojórquez E, González-Peláez SS (2022) The last geoduck:The experience of geoduck clam fishery management in the Mexican Pacific Ocean. Mar Policy 143:105145

McClenachan L, Ferretti F, Baum JK (2012) From archives to conservation: why historical data are needed to set baselines for marine animals and ecosystems. Conservation Letters 5:349–359

McLain R, Lawry S, Ojanen M (2018) Fisheries’ property regimes and environmental outcomes:A realist synthesis review. World Dev 102:213–227

Mellink E, Orozco-Meyer A (2006) Abundance, distribution, and residence of bottlenose dolphins (*Tursiops truncatus*) in the Bahia San Jorge Area, Northern Gulf of California, Mexico. Aquat Mamm 32:133–139

Mitchell DR, Foster MS (2000) Hohokam Shell Middens along the Sea of Cortez, Puerto Peñasco, Sonora, Mexico. J Field Archaeol 27:27–41

Mitchell DR, Huckleberry G, Rowell K, Dettman DL (2015) Coastal Adaptations During the Archaic Period in the Northern Sea of Cortez, Mexico. The Journal of Island and Coastal Archaeology 10:28–51

Moreno-Báez M, Cudney-Bueno R, Orr BJ, et al (2012) Integrating the spatial and temporal dimensions of fishing activities for management in the Northern Gulf of California, Mexico. Ocean Coast Manag 55:111–127

Morzaria-Luna HN, Castillo-López A, Danemann GD, Turk-Boyer PJ (2014a) Conservation strategies for coastal wetlands in the Gulf of California, Mexico. Wetlands Ecol Manage 22:267–288

Morzaria-Luna H, Turk-Boyer PJ, Rosemartin A, Camacho-Ibar VF (2014b) Vulnerability to climate change of hypersaline salt marshes in the Northern Gulf of California. Ocean Coast Manag 93:37–50

Munguía-Vega A, Torre J, Turk-Boyer PJ, et al (2015) PANGAS: An Interdisciplinary Ecosystem-Based Research Framework for Small-Scale Fisheries in the Northern Gulf of California. J Southwest 57:337–390

Munro Palacio G (2007) Breve historia de Puerto Peñasco. De Cierto Mar Editores, Puerto Peñasco, Sonora

Naranjo-Madrigal H, van Putten I, Norman-López A (2015) Understanding socio-ecological drivers of spatial allocation choice in a multi-species artisanal fishery: A Bayesian network modeling approach. Mar Policy 62:102–115

Oksanen J, Blanchet G, Kindt R, et al (2010) vegan: Community Ecology Package

Olsson P, Folke C (2001) Local ecological knowledge and institutional dynamics for ecosystem management: A study of Lake Racken watershed, Sweden. Ecosystems 4:85–104

Ortlieb L (1991) Quaternary Vertical Movements Along the Coasts of Baja California and Sonora: Chapter 22: Part III. Regional Geophysics and Geology. In: Dauphin JP, Simoneit BT (eds) The Gulf and Peninsular Province of the Californias. AAPG Special Volumes, pp 447–480

Palomares M, Pauly D (2014) SeaLifeBase. World Wide Web electronic publication. www.sealifebase.org, version (11/2014). http://www.sealifebase.ca/

Pauly D (1995) Anecdotes and the shifting baseline syndrome of fisheries. Trends Ecol Evol 10:430

Pauly D, Christensen V (1995) Primary production required to sustain global fisheries. Nature 374:255–257

Pauly D, Christensen V, Dalsgaard J, et al (1998a) Fishing Down Marine Food Webs. Science 279:860–863

Pauly D, Trites AW, Capuli E, Christensen V (1998b) Diet composition and trophic levels of marine mammals. ICES J Mar Sci 55:467–481

Previero M, Gasalla MA (2018) Mapping fishing grounds, resource and fleet patterns to enhance management units in data-poor fisheries: The case of snappers and groupers in the Abrolhos Bank coral-reefs (South Atlantic). Ocean Coast Manag 154:83–95

R Development Core Team (2021) R: A language and environment for statistical computing. Ver 4.1.1. R Foundation for Statistical Computing, Vienna, Austria

Rick TC, Reeder-Myers LA, Hofman CA, et al (2016) Millennial-scale sustainability of the Chesapeake Bay Native American oyster fishery. Proc Natl Acad Sci U S A 113:6568–6573

Robertson HA, McGee TK (2003) Applying local knowledge: the contribution of oral history to wetland rehabilitation at Kanyapella Basin, Australia. J Environ Manage 69:275–287

Rodríguez-Quiroz G, Aragón-Noriega A, Valenzuela-Quiñónez W, Esparza-Leal HM (2010) Pesca artesanal en las zonas de conservación del Alto Golfo de California. Rev Biol Mar Oceanogr 45:89–98

Rose KA, Roth BM, Smith EP (2009) Skill assessment of spatial maps for oceanographic modeling. J Mar Syst 76:34–48

Rubio-Cisneros NT, Aburto-Oropeza O, Jackson J, Ezcurra E (2017) Coastal Exploitation Throughout Marismas Nacionales Wetlands in Northwest Mexico. Tropical Conservation Science 10:1940082917697261

Sáenz-Arroyo A, Revollo-Fernández D (2016) Local ecological knowledge concurs with fishing statistics: An example from the abalone fishery in Baja California, Mexico. Mar Policy 71:217–221

Sáenz-Arroyo A, Roberts CM, Torre J, et al (2005a) Rapidly shifting environmental baselines among fishers of the Gulf of California. Proceedings of the Royal Society of London, Series B: Biological Sciences 272:1957–1962

Sáenz-Arroyo A, Roberts CM, Torre J, Carino-Olvera M (2005b) Using fishers’ anecdotes, naturalists’ observations and grey literature to reassess marine species at risk: the case of the Gulf grouper in the Gulf of California, Mexico. Fish Fish 6:121–133

Sagarin RD, Gilly WF, Baxter CH, et al (2008) Remembering the Gulf: changes to the marine communities of the Sea of Cortez since the Steinbeck and Ricketts expedition of 1940. Front Ecol Environ 6:372–379

Sala E, Aburto-Oropeza O, Reza M, et al (2004) Fishing down coastal food webs in the Gulf of California. Fisheries 29:19–25

Sawatzky DL, Raines GL, Bonham-Carter GF, Looney CG (2009) Spatial Data Modeller (SDM):ArcMAP 9.3 geoprocessing tools for spatial data modelling using weights of evidence, logistic regression, fuzzy logic and neural networks

Scheffer M, Carpenter S, Young B de (2005) Cascading effects of overfishing marine systems. Trends Ecol Evol 20:579–581

Seminoff JA, Nichols WJ (2006) Sea turtles of the Alto Golfo. A struggle for survival. In: Felger RS, Broyles B (eds). University of Utah Press, Salt Lake City, Utah, pp 505–518

Serra-Pereira B, Erzini K, Maia C, Figueiredo I (2014) Identification of potential essential fish habitats for skates based on fishers’ knowledge. Environ Manage 53:985–998

Sierra R, Chamberlain J (1999) Mapping and monitoring of coastal wetlands in Sonora, Mexico:a multi-national approach. Department of Geography. Arizona State University

Silvano RAM, Valbo-Jørgensen J (2008) Beyond fishermen’s tales: contributions of fishers’ local ecological knowledge to fish ecology and fisheries management. Environ Dev Sustainability 10:657

Soga M, Gaston KJ (2018) Shifting baseline syndrome: causes, consequences, and implications. Front Ecol Environ 16:222–230

Spackeen J (2009) Analysis of food web dynamics in the Northern Gulf of California using stable isotopes

Thomson DA, Mead AR, Schreiber JF (1968) Probable environmental impact of heated brine effluents from a nuclear desalination plant on the Northern Gulf of California. University of Arizona, Marine Science Committee. U.S. Dept. of Interior. Office of Saline Water, Research and Development, Tucson, AZ

Turk-Boyer PJ (2008) Integrating science and community participation: Conservation and management of coastal wetlands in the Northern Gulf of California. The David and Lucile Packard Foundation

Turk-Boyer PJ, Morzaria-Luna H, Martínez I, et al (2014a) Ecosystem-based fisheries management of a biological corridor along Northern Sonora Coastline. In: Amezcua F, Bellgraph B (eds) Fisheries Management of Mexican and Central American Estuaries. Springer Verlag, pp 155–180

Turk-Boyer P, Peña-Bonilla H, Morzaria-Luna HN, et al (2014b) Wetland conservation in Northern Sonora, Mexico: Legal tools and active communities. In: Fisheries Management of Mexican and Central American Estuaries. Springer, pp 181–204

Turvey ST, Barrett LA, Yujiang H, et al (2010) Rapidly shifting baselines in Yangtze fishing communities and local memory of extinct species. Conserv Biol 24:778–787

Valdes-Casillas C, Carrillo-Guerrero Y, Zamora-Arroyo F, et al (1999) Mapping and management of coastal wetlands of Puerto Peñasco, Sonora: A multinacional project. Center for Conservation of Natural Resources (CECARENA), Instituto Tecnológico y de Estudios Superiores de Monterrey -- Campus Guaymas (ITESM-CG); Pronatura. Arizona State University, Sonora

Van Dyke E, Wasson K (2005) Historical ecology of a central California estuary: 150 years of habitat change. 28:173–189

Venables WN, Ripley BD (2002) Modern applied statistics with S. Springer, New York

Wanyonyi IN, Macharia D, Heenan A, Mangi SC (2018) Using participatory methods to assess data poor migrant fisheries in Kenya. Hum Dimensions Wildl 23:569–586

Wilson KA, Westphal MI, Possingham HP, Elith J (2005) Sensitivity of conservation planning to different approaches to using predicted species distribution data. Biol Conserv 122:99–112

Zadeh LA (1965) Fuzzy sets. Information and Control 8:338–353

Zedler JB, Kercher S (2005) Wetland resources: Status, trends, ecosystem services, and restorability. Annu Rev Environ Resour 30:39–74

Zu Ermgassen PSE, Spalding MD, Blake B, et al (2012) Historical ecology with real numbers:past and present extent and biomass of an imperilled estuarine habitat. Proc Biol Sci 279:3393–3400

