## Supplementary information for "Historical use of coastal wetlands by small-scale fisheries in the Northern Gulf of California"

### Historical patterns of commercial species abundance and fishing areas in coastal wetlands of the Northern Gulf of California

Hem Nalini Morzaria-Luna[1,2], Mabilia Urquidi[1,3], Gabriela Cruz-Piñón[4], Jośe Manuel Dorantes Hernández[4,6], Paloma A. Valdivia Jiménez[1], Ángeles Sánchez Cruz[1], Ilse Martínez[5]

1. CEDO Intercultural, PO Box 44208, Tucson, AZ 8573, USA. / Edif. Agustín Cortés, s/n. Puerto Peñasco, Sonora CP 83550, México.
3. Current address: Posgrado en Ciencias Marinas y Costeras. Universidad Autónoma de Baja California Sur. Carretera al Sur Km 5.5, Apartado Postal 19-B, C.P. 23080. La Paz, Baja California Sur.,
4. Departamento de Ciencias Marinas y Costeras. Universidad Autónoma de Baja California Sur. Carretera al Sur Km 5.5, Apartado Postal 19-B, C.P. 23080. La Paz, Baja California Sur. GCP,
5. Department of Environmental Studies. University of Victoria. David Turpin Building, B243. Victoria BC Canada,.
6. Current address: El Colegio de la Frontera Sur (ECOSUR). Av. Rancho Polígono 2-A Ciudad Industrial Lerma, San Francisco de Campeche, Campeche, México.

#### Data repository
https://knb.ecoinformatics.org/view/doi%3A10.5063%2FF12R3Q48

**Text S1. Informed consent form for expert users.**

**RESEARCH CONSENT FORM**

**FOR QUESTIONS ABOUT THE STUDY, CONTACT:** Dr. Hem Nalini Morzaria Luna. CEDO Intercultural.. (XXX) XXX XXXX

**DESCRIPTION:** You are invited to participate in a research study on historical ecology of the Northern Gulf. The purpose of this survey is to assess the knowledge and experience of expert users of Northern Gulf Wetlands, along the coast of Sonora. We expect to understand the rate of change in different sites and to document the impacts the wetlands have suffered in the last century. You will be asked to answer a survey, the survey will be conducted via telephone or in person. Personal and phone interviews will be taped in electronic format. The files from the interviews will be kept in a computer only accessible to CEDO researchers.

**RISKS AND BENEFITS:** There are no risks associated with this study. The benefits which may reasonably be expected to result from this study are contributing to further understanding ecology of Northern Gulf wetlands. We cannot and do not guarantee or promise that you will receive any benefits from this study.

**TIME INVOLVEMENT:** Your participation in this study will take approximately 1 hour.

**PAYMENTS:** You will not receive a compensation from participating in this study. SUBJECT'S RIGHTS: If you have read this form and have decided to participate in this project, please understand your participation is voluntary and you have the right to withdraw your consent or discontinue participation at any time without penalty or loss of benefits to which you are otherwise entitled. You have the right to refuse to answer particular questions. If you agree, your identity will remain confidential in all written data resulting from the study. With your specific consent, we will attribute specific quotes or comments.

**Contact Information:**
**Questions, Concerns, or Complaints:** If you have any questions, concerns or complaints about this research study, its procedures, risks and benefits, or alternative courses of treatment, you should ask the Protocol Director, Hem Nalini Morzaria Luna. You may contact him/her now or later at. (XXX) XXX XXXX

**Independent Contact:** If you are not satisfied with how this study is being conducted, or if you have any concerns, complaints, or general questions about the research or your rights as a participant, please contact CEDO's subdirector, Alejandro Castillo Lopez to speak to someone independent of the research team at (XXX) XXX XXXX

**Text S2. Consent form for resource users.** Translated from the original in Spanish.

Interview Authorization
Good morning/afternoon. My name is… I work for the Intercultural Center for the Study of Deserts and Oceans (CEDO) on two projects focusing on the history of estuaries in the North Gulf and the modeling of fish populations (Atlantis).

The purpose of the project is to understand fishing activities in the wetlands of the northern Gulf of California and how these areas have changed over time. As part of our project, we are interviewing resource users in the region to allow us to learn a little more about which estuaries are important for fishing, the species that breed in the estuaries, and what changes have been observed in the estuaries, in order to understand the ecology of the estuaries and their history.

We would like to invite you to voluntarily participate in this project. We invite you because you are an experienced fisher in the northern Gulf of California. Your participation involves an interview on the topics I just mentioned. The interviews will be held at a location of your convenience and will last approximately two hours. There are no risks associated with your participation in these interviews and no direct benefits are expected from your participation. Likewise, there is no cost to you except your time, but you will not be compensated in monetary form for your participation.

The information you provide us is strictly confidential and will only be used for research purposes. During the interviews, we will take notes to help us review what was said. Only the principal investigator will have access to your name and the information you provide. To maintain your anonymity, your name will not be featured in any reports resulting from this project.

Your participation in this study is completely voluntary and you may stop participating at any time if you feel it is necessary. Any questions you have will be answered and you can decide which questions you want or can answer. However, all your opinions are very important to us and we hope that you will want to participate.

By participating in the interviews, you are giving us permission to use your information for research purposes.

**Text S3. Historical ecology questionnaire applied to oyster producers.** Translated from the original Spanish.

*Before starting, the interviewer must provide the interviewee with the information contained in the informed consent document.*

| | |
|---|---|
| Community where the interview is conducted | Interview time (24 hr format) |
| Interviewer's name | Date (month/day/year) |
| Survey number<br>O - | Do you consent to being interviewed?<br>Yes No |

**INTERVIEWEE'S GENERAL INFORMATION**

| | |
|---|---|
| 1. Name/Nickname | 2. Age |
| 3. City of residence and residence time | 4A.Do you actively work in aquaculture?<br>Yes No |
| 4B.Besides being an oyster producer, have you ever been a fisher ? Yes No<br>Are you an active fisher? Yes No | 5. What kind of fishing?<br>Recreational Commercial Artisanal |
| 6.A How many years were you a fisher and in what years??<br>Age (or year) at which you started fishing _______<br>At what age did you stop fishing? ____________<br>Why did he become an oyster producer? | 7. Number of years as an oyster producer??<br>In what year did you start?<br>(If necessary, please estimate the interval):<br>(0-5) (5-10) (10-20) (20-30) (30-40) (40+)<br>Cooperative name: |
| 8 Are any of your children / father an oyster producer or fisher?<br>Yes No Oyster producer____ Fisher___<br>In what community? | 9 What is / was the main fishing art or techniques?<br>10A What is / was the main species fished? |
| 10B. What is/was the main species they cultivated? | 10. What other species have you cultivated? |
| 11. On average, how many months per year is dedicated to aquaculture? And how many months per year to fishing?<br>12. Have you fished inside some Estuary, Salina, or Bay?<br>___Have you fished to sell the products<br>___Have you fished for subsistence (you or your family ate what they caught) | 12. In which estuary do you currently have crops or which was the last one you cultivated? |

**NOTE:IF THE OYSTER PRODUCER IS ALSO A FISHER, IT IS NECESSARY TO FILL THE APPENDIX**

**GENERAL USE OF ESTUARIES**

13. Can you tell me which estuary you know and have done the following activities?
[Show general map of the area with the estuary marked in color or the toponymy map]

| Estero | Aquaculture | Fishing | Recreation | Another |
|---|---|---|---|---|
| La Salina | | | | |
| San Francisquito | | | | |
| La Cinita | | | | |

| |
|---|
| Estero Almejas |
| La Pinta |
| Morua |
| La Cholla |
| Cerro Prieto |
| Las Lisas |
| San Judas |
| Bahía Adair |
| Other |

*I will ask you to help me draw the estuary with which you are most familiar .. Draw it as it was the first time I arrived*

**CAPTURED SPECIES**

14. Next, I will read a list of species.
Can you identify which species breed within estuaries? Indicate which ones breed within them, and which ones did but not anymore. Do you know fishing areas for these species?
*NOTE: If the interviewee uses a different common name, write it underneath the species name. Estimate the date when referring to historical data.*

| Species | Estuary | Season (present/historical) | Sustenance fishing (S) o commercial (C) | Breeding zone |
|---|---|---|---|---|
| Clam | | | | |
| Blue shrimp | | | | |
| Black and pink murex snail | | | | |
| Conch | | | | |
| Brown and blue crab | | | | |
| Oysters | | | | |
| Whiptail stingray | | | | |
| California butterfly ray | | | | |
| Guitarfish | | | | |
| Shortfin weakfish | | | | |
| Spotted sand bass | | | | |
| Mullet | | | | |
| Pompano | | | | |
| Mackerel | | | | |
| Mojarra | | | | |
| Flatfish | | | | |

15. I will now show you maps of the estuaries and I will ask questions related to breeding and nursery areas, fishing areas, and other important areas.
*[Maps of each estuary, one estuary per page.The interviewee should be encouraged to be as specific in fishing areas as possible. Focus on species and species estuaries mentioned by the interviewee]*

Breeding zone, Fishing areas, Areas where they cultivate oysters, Tourism (areas where tourists arrive or where they take tourists in case they do), Bird nesting (Examples: Gallito de Mar, Herons, Seagulls)

16. For the three main species you select present within estuaries, do you know when the breeding occurred? and fishing?

| Breeding/fishing | Species | Jan | Feb | Mar | April | May | Jun | Jul | Aug | Sep | Oct | Nov | Dec |
|---|---|---|---|---|---|---|---|---|---|---|---|---|---|
| Breeding | | | | | | | | | | | | | |
| Fishing | | | | | | | | | | | | | |
| Breeding | | | | | | | | | | | | | |
| Fishing | | | | | | | | | | | | | |
| Breeding | | | | | | | | | | | | | |
| Fishing | | | | | | | | | | | | | |

17. While you have been an oyster producer, have you observed changes in the breeding areas or when the species getting inside in the estuary? [focus on the three previous species]
How often was the change? In what years was this change observed?
18. What species did you observe breeding within the estuaries of the region (in the time you have worked in them)?
18 a. Has the abundance of the species in the estuary changed (in whole/ live weight in kg), has it increased or decreased compared to the current time?.
18. b. During the time you have fished, where there changes in species raised in the estuaries? Changes in areas or size? Indicate what species.
18. c. Was there any extreme reduction or did any species disappear?
These changes were noticeable in the following decades:

| Species | Increase / Decrease (+ / 0 / -) and write the year of sudden event | | | | | | Changes in Breeding Zone (S/N) | Extreme reduction or disappeara nce (S/N) |
|---|---|---|---|---|---|---|---|---|
| | 2,000 | 1990 | 1980 | 1970 | 1960 | 1950 | | |
| Clam | | | | | | | | |
| Blue shrimp | | | | | | | | |
| Black and pink murex snail | | | | | | | | |
| Conch | | | | | | | | |
| Brown and blue crab (jaiba) | | | | | | | | |
| Oysters (ostiones) | | | | | | | | |

| |
|---|
| Whiptail stingray (manta arenera) |
| California butterfly ray (manta mariposa) |

| Species | Increase/Decrease (+/0/-) and score year of subdue event | | | | | | Changes in breeding area (S/N) | Extreme Reduction or Disappearance (R/D) |
|---|---|---|---|---|---|---|---|---|
| | 2000 | 1990 | 1980 | 1970 | 1960 | 1950 | | |
| Guitarfish | | | | | | | | |
| Pufferfish | | | | | | | | |
| Corvina de orilla | | | | | | | | |
| Cabrilla arenera | | | | | | | | |
| Lisa | | | | | | | | |
| Branches | | | | | | | | |
| Sierra | | | | | | | | |
| Mojarras | | | | | | | | |
| Flatfish<br>Seabirds<br>Other | | | | | | | | |

19.What do you think are the main causes or responsible for these changes?

**Estuaries**

20. Have you noticed the following changes in the estuaries of the area over time? What kind of changes and in what areas? *[The following table help to guide the question, but the surveyed should be encouraged to answer the question freely and tell stories about specific sites if possible]*

| Conditions | Good Changes | Bad Changes | Not change | Cause of change |
|---|---|---|---|---|
| Area covered by salted grass or plants | | | | |
| Animals or plants (number or types) | | | | |
| Shape and type of channels | | | | |
| Oyster production | | | | |
| Tourism developments | | | | |

Erosion dunes

Some other change

21. Which estuary do you think has suffered bigger impacts and of what type?
22.How do you think these impacts affect the fishing production of the estuaries and outside the estuaries?
23. What are threats exist today for estuaries? Do you think that these threats have changed over time?

| a) 2000 to date | b) 1990 | c) 1980 | d) 1970 | d) 1960 | e) before 1950 |
|---|---|---|---|---|---|

24. Have you reported wetland destruction to the authorities? Write to which authority
Would you be interested in learning how to do it?
25. Do you consider that these estuaries have some federal protection level that prohibits their destruction?
26.If you could protect only one of the estuaries mentioned at the beginning, which one would you choose?

**APPENDIX A. SPECIES OUTSIDE THE ESTUARIES**

27. Change in abundance: For each decade, indicate whether the relative abundance of the fish or animal, such as high abundance (+), medium abundance (0), or low abundance (-).
Was there an extreme increase in fish price? If so, indicate in what year.
Has there been a reduction in the number of animals or have they disappeared? Yes or no
Has the size of the animals been reduced? Yes or no

| | | **Change of Abundance (-/0/+)** | | | | | | **Change in price** | **Reduction** | **Smaller fish** |
|---|---|---|---|---|---|---|---|---|---|---|
| **Species** | | **1950** | **1960** | **1970** | **1980** | **1990** | **2000** | **Year** | **(S/N)** | **(S/N)** |
| Gulf coney | *Epinephelus acanthistius* | | | | | | | | | |
| Goldspotted sandbass | *Paralabrax auroguttatus* | | | | | | | | | |
| Leopard grouper | *Mytoperca rosacea* | | | | | | | | | |
| Gulf grouper | *Myctoperca jordani* | | | | | | | | | |
| Sea bass | Serranidos | | | | | | | | | |
| Yellow snapper | *Lutjanus argentiventris* | | | | | | | | | |
| Pargo coconaco | *Hoplopagrus guentherii* | | | | | | | | | |
| Snapper | Lutjanidae | | | | | | | | | |
| Milkfish | Sciaenidae | | | | | | | | | |
| Chicharro | Haemulidae | | | | | | | | | |
| Parrot fish | Scaridae | | | | | | | | | |
| Surgeon fish | Acanthuridae | | | | | | | | | |
| Large reef fish | Fish like pigs | | | | | | | | | |
| Small reef fish | Fish such as boetetes and gobidos | | | | | | | | | |
| Angel shark | *Squatina californica* | | | | | | | | | |

| | |
|---|---|
| Tripa | Mustelus spp. |
| Large pelagic sharks | Selachimorpha |

| | | **Change of Abundance (-/0/+)** | | | | | | **Change in price** | **Reduction** | **Smaller fish** |
|---|---|---|---|---|---|---|---|---|---|---|
| **Species** | | **1950** | **1960** | **1970** | **1980** | **1990** | **2000** | **Year** | **(S/N)** | **(S/N)** |
| Guitar | Rhinobatidae | | | | | | | | | |
| Stripes, blankets and sharks | Butterfly and Diamond Blanket | | | | | | | | | |
| Flatfish | Pleuronectidae, Bothidae, Paralichthyidae | | | | | | | | | |
| Mojarra | Gerreidae | | | | | | | | | |
| Scorpion fish | Scorpaenidae, Triglidae | | | | | | | | | |
| Deep fish | Myctophidae | | | | | | | | | |
| Totoaba | Totoaba macdonaldi | | | | | | | | | |
| Major pelagics | e.g.,Sphyraenidae, Carangidae, Istiophoridae | | | | | | | | | |
| Sierra | Scombridae | | | | | | | | | |
| Hake | Merlucciidae | | | | | | | | | |
| Anchovies, sardines | Minor pelagics | | | | | | | | | |
| Jaiba and lobster | Callinectes bellicosus, C.arcuatus, OrderDecapoda | | | | | | | | | |
| Callo and clams | e.g.,Spondylus calcifer,S. princeps, Pteria sterna | | | | | | | | | |
| Clams, mussels, oysters | Bivalvia | | | | | | | | | |
| Shrimp | Penaeidae | | | | | | | | | |
| Sea cucumber | Holothuroidea | | | | | | | | | |
| Sea urchins | Echinodermata | | | | | | | | | |
| Snail | Gastropoda | | | | | | | | | |

| | |
|---|---|
| Squid | Teuthoidea |
| Whale (mysticets) | Fin Whale, Humpback Blue Ballen |
| Whale (odontocetes) | Orcas, Sperm Whales Pilot Whales |
| Dolphin | *Delphinus capensis, D. delphis*,Tursiops truncatus, Lagenorhynchus obliquidens |
| Vaquita porpoise | *Phocoena sinus* |
| Sea lions and seals | Pinnipeds |
| Turtles | Testudines |
| Seabirds | Seagulls, Pelicans, Cormorants |

Interviewer's remarks:

**Text S4. Historical ecology questionnaire applied to fishers.** Translated from the original Spanish.

***Before starting, the interviewer must provide the interviewee with the information contained in the informed consent document.***

| | |
|---|---|
| Community where the interview takes place | Interview time (24 hr format)<br>From : to : |
| Interviewer's name | Date (month/day/year) |
| Survey number | Do you agree to be interviewed?<br>Yes No |

**INTERVIEWEE'S GENERAL INFORMATION**

| | |
|---|---|
| 1. Name/Nickname | 2. Age |
| 3. City of residence and residence time | 4. Are you an active fisher? Yes No |
| 5 What kind of fishing do you do or did (can be more than one)?<br>Recreational Artisanal Commercial | 6. How many years were you a fisher? |
| 7. In what year did you start fishing?<br>(If necessary, please estimate the interval):<br>(0-5) (5-10) (10-20) (20-30) (30-40) (40+) | 8.Any of your children / father is a fisher??<br>Yes No<br>In what community |
| 9.What is / was your main fishing gear? | 10. What is/was your main fishing species? |
| 11.On average, how many months per year did you spend fishing?? | 12.Have you fished in some estuary, Sslina, or bay??<br><br>Yes No<br><br>____Have you fished to sell the products?<br>____Have you fished for your own use (you or your family ate what they caught) |
| Comments: | |

**GENERAL USE OF ESTUARIES**

**13 Can you tell me in which estuaries you have fished and in which you have done other activities besides fishing?**
***[Show general map of the area with the estuaries marked in color or the toponymy map]***

| Estero | Fishing | Recreation | Aquaculture | Other |
|---|---|---|---|---|
| La Salina | | | | |
| San Francisquito | | | | |
| La Cinita | | | | |
| Almejas | | | | |
| La Pinta | | | | |

| |
|---|
| Morua |
| La Cholla |
| Cerro Prieto |
| Las Lisas |
| San Judas |
| Bahía Adair |
| Other |

***Now I am going to ask you to help me draw the estuary that you have visited the most or where you have fished the most. Draw it as it was when you started fishing.***

**TARGET SPECIES**

**14. I will read you a list of species. Please indicate which of these species you fish or have fished in the estuaries and with what fishing gear.**

**Can you identify which species breed or use estuaries as nursery sites? Indicate which ones were raised, but not anymore, and which ones are still raised**

***NOTE: If the surveyed uses a different common name, write it down under the species box.***
***Estimate the date when it is historical*** **data**

| Species | | Estuary | Time (present/historical) | Commercial/Sustenance fishing | Breeding area |
|---|---|---|---|---|---|
| Clam | | | | | |
| Blue shrimp | | | | | |
| Black and pink murex snail | | | | | |
| Conch | | | | | |
| Brown and blue Crab | | | | | |
| Oysters | | | | | |
| Whiptail stingray | | | | | |
| California butterfly ray | | | | | |
| Guitarfish | | | | | |
| Pufferfish | | | | | |
| Shortfin weakfish | | | | | |
| Spotted sand bass | | | | | |
| Mullet | | | | | |
| Pampanos | | | | | |
| Sierra | | | | | |
| Mojarra | | | | | |

| Flatfish | | | | | |
|---|---|---|---|---|---|
| Species | | Estuary | Time (present/historical) | Commercial/Sustenance fishing | Breeding area |
| Other<br><br>Other<br><br>Other | | | | | |

**15. Next, I will show you maps of the estuaries and I will ask you questions related to the areas in which you fish, the sites that you consider are important for the maintenance of this fishery, and other important areas.**

***[Maps of each estuary, a estuary per page. The surveyed will be encouraged to be as specific in fishing areas as possible. Focus on the species and estuaries mentioned by the interviewee]***

**Ask the fisherman to identify according to their knowledge (past and present): Breeding areas, Fishing areas, Tourism (areas where tourists arrive or where they take tourists in case they do), Bird nesting (Examples: Gallito de Mar, Herons, Seagulls)**

**16 In which months of the year do you fish (-ed) the most important species? [Focus on three species they extracted in more estuaries]**

| Breeding/fishing | Species | Jan | Feb | Mar | April | May | Jun | Jul | Aug | Sep | Oct | Nov | Dec |
|---|---|---|---|---|---|---|---|---|---|---|---|---|---|
| Breeding | | | | | | | | | | | | | |
| Fishing | | | | | | | | | | | | | |
| Breeding | | | | | | | | | | | | | |
| Fishing | | | | | | | | | | | | | |
| Breeding | | | | | | | | | | | | | |
| Fishing | | | | | | | | | | | | | |

**17. Along the time you were a fisherman, did you observe changes in these 3 important species during fishing times in the estuary? How? (For example, did they get longer, shorter or did they occur in another month than usual?)**

**How often was the change?**

**In what years did you observe this change?**

**18. What species did you observe that use the estuaries in the region as breeding or nursery areas (over the time you fished)?**

**18 a. Has target species abundance (in whole weight / live in kg) increased or decreased compared to the current time?**

**18 b. Since you started fishing have you noticed changes in species that grow within estuaries? Changes in area or size. Indicate what species.**

**18 c. Did you observe extreme reductions or the disappearance of any species?**

**Indicate which of these changes were noticeable in the following time periods:**

| Species | Increase/Decrease (-/0/+) | | | | | | Changes in breeding area (S/N) | Extreme reduction or disappearance (S/N) |
|---|---|---|---|---|---|---|---|---|
| | 2000 | 1990 | 1980 | 1970 | 1960 | 1950 | | |
| Clam | | | | | | | | |
| Blue shrimp | | | | | | | | |
| Black and pink murex snail | | | | | | | | |
| Conch | | | | | | | | |
| Brown and blue crab | | | | | | | | |
| Oysters | | | | | | | | |
| Whiptail stingray | | | | | | | | |
| California butterfly ray | | | | | | | | |
| Guitarfish | | | | | | | | |
| Pufferfish | | | | | | | | |
| Shortfin weakfish | | | | | | | | |
| Spotted sand bass | | | | | | | | |
| Mullet | | | | | | | | |
| Pompano | | | | | | | | |
| Mackerel | | | | | | | | |
| Mojarra | | | | | | | | |
| Flatfish<br>Other | | | | | | | | |

##### 19. What do you think are the main causes or or drivers of these changes?

#### Estuaries

**20. Have you noticed changes in estuaries areas over time? What kind of changes and in what areas?**

| Conditions | Positive changes | Negative changes | No change | Cause of change |
|---|---|---|---|---|

| |
|---|
| **Area covered by salt grass or plants** |
| **Animals or plants (number or types)** |
| **Shape and type of channels** |
| **Oyster production** |
| **Nearby developments (constructions)** |
| **Dune erosion** |

**21. Which estuary do you think has suffered bigger impacts and of what type?**

**22. How do you think these impacts affect the fisheries production within and outside estuaries?**

**23. What are the main threats today for estuaries? Do you think that these threats have changed over time?**

| a) 2000 until current time | b) 1990 | c) 1980 | d) 1970 | d) 1960 | e) before 1950 |
|---|---|---|---|---|---|

**24. Have you reported wetland destruction to the corresponding authorities? Indicate which authority**

**Would you be interested in learning how to report environmental damage?**

**25. Do you think estuaries should have federal protection to prohibit their destruction?**

**26. If you could protect only one of the estuaries mentioned at the beginning of this survey, which one would you choose?**

**SPECIES OUTSIDE ESTUARIES**

**27. Change in abundance: For each decade, indicate changes in the relative abundance of the species, as high abundance (+), medium abundance (0), and low abundance (-).**

**Was there an extreme increase in fish price (did a market develop?). If so, indicate in what year.**

**Has there been an extreme reduction in species abundance or local extictions? Yes or no.**

**Has the size of the animals been reduced? Yes or no**

| | | Change of Abundance (-/0/+) | Change in price | Reduction | Smaller fish |
|---|---|---|---|---|---|

| Species | | 1950 | 1960 | 1970 | 1980 | 1990 | 2000 | Year | (S/N) | (S/N) |
|---|---|---|---|---|---|---|---|---|---|---|
| Gulf coney | *Epinephelus acanthistius* | | | | | | | | | |
| Goldspotted sandbass | *Paralabrax auroguttatus; Paralabrax parrot* | | | | | | | | | |
| Leopard grouper | *Mytoperca rosacea* | | | | | | | | | |
| Gulf grouper | *Myctoperca jordani* | | | | | | | | | |
| Groupers | Serranidae | | | | | | | | | |
| Yellow snapper | *Lutjanus argentiventris* | | | | | | | | | |
| Mexican barred snapper | *Hoplopagrus guentherii* | | | | | | | | | |
| Snapper | Lutjanidae | | | | | | | | | |
| Milkfish | Sciaenidae | | | | | | | | | |
| Scad | Carrangidae | | | | | | | | | |
| Parrot fish | Scaridae | | | | | | | | | |
| Surgeonfish | Acanthuridae | | | | | | | | | |
| Larger reef fish | Fish like triggerfish | | | | | | | | | |
| Smaller reef fish | Fish such as pufferfish and gobiers | | | | | | | | | |
| Angel shark | *Squatina californica* | | | | | | | | | |
| Smoothhound shark | Mustelus spp. | | | | | | | | | |
| Large pelagic sharks | Selachimorpha | | | | | | | | | |
| Guitarfish | Rhinobatidae | | | | | | | | | |
| Rays, mantarrays and sharks | Butterfly and Diamond ray | | | | | | | | | |
| Flatfish | Pleuronectidae, Bothidae, Paralichthyidae | | | | | | | | | |
| Mojarra | Gerreidae | | | | | | | | | |
| Scorpion fish | Scorpaenidae, Triglidae | | | | | | | | | |
| Demersal fish | Myctophidae | | | | | | | | | |
| Totoaba | *Totoaba macdonaldi* | | | | | | | | | |
| Large pelagics | Sphyraenidae, Carangidae, Istiophoridae | | | | | | | | | |
| Sierra | Scombridae | | | | | | | | | |
| Hake | Merlucciidae | | | | | | | | | |
| Anchovies, sardines | Small pelagics | | | | | | | | | |
| Blue and brown crabs, lobster | *Callinectes bellicosus, C. arcuatus*, Order Decapoda | | | | | | | | | |

| | |
|---|---|
| Scallops and clams | *Spondylus calcifer, S. princeps, Pteria sterna* |
| Clams, mussels, oysters | Bivalvia |
| Shrimp | Penaeidae |
| Sea cucumber | Holothuroidea |
| Sea urchins | Echinodermata |
| Snails | Gastropoda |
| Squid | Teuthoidea |
| Whales (mysticetes) | Fin Whale, Humpback Blue Ballen |
| Whale (odontocetes) | Orcas, Sperm whales, Pilot whales |
| Dolphins | *Delphinus capensis, D. delphis, Tursiops truncatus, Lagenorhynchus obliquidens* |
| Vaquita porpoise | Phocoena sinus |
| Sea lions and seals | Pinnipeds |
| Sea turtles | Testudines |
| Seabirds | Seagulls, Pelicans, Cormorants |

**Interviewer's remarks:**

**Text S5. Historical ecology interview questions applied to expert users.**

The purpose of this survey is to assess the knowledge and experience of expert users in Northern Gulf Wetlands. We expect to understand the rate of change in different sites and to document the impacts the wetlands have suffered. Before you begin, please read the attached informed consent form. The interviewee should understand this information. Northern Gulf wetlands refer to the wetlands in the coast of Sonora between El Borrascoso and San Francisquito, south of Bahia San Jorge.

| |
|---|
| Interviewer/ Transcribed by: |
| Date (m/d/y): |
| Time: |
| Interview number: |
| Interview duration: |
| WAV File name: |

#### GENERAL INFORMATION

| | |
|---|---|
| 1. Name: | 2. Institution of affiliation: |
| 3. Sex: | 4. Other institutions of affiliation (please list in last ten years): |
| 5. Age (for statistical purposes): | |
| 6. Position: | 7. Main research interest: |
| 8. In what year did you first visit the Gulf of California? | 9. How many times have you visited the Gulf:<br>Bin responses as 0, 1-5, >5, >10 times<br>a) Before 1970<br>b) Between 1970-1990<br>c) Between 1990-2000<br>d) After 2000- 2008 |
| 10. Main purpose of your visits to the Gulf of California<br>a. Research<br>b. Teaching<br>c. Thesis (M.S. or Ph.D.) | 11. Do you have any current research projects/interests in the Gulf of California?<br>YES NO<br><br>Project title: |

#### GENERAL WETLAND QUESTIONS

12. ¿Have you visited any coastal wetland, Salina, or bay in the Northern Gulf? YES/NO

13. a. Can you tell me which wetlands have you visited? And what activities you have carried out? Explain nature of each activity as indicated

b. When was the last time you visited any of these coastal wetlands?

Northern Gulf wetlands refer to the wetlands in the coast of Sonora between El Borrascoso and San Francisquito, south of Bahia San Jorge

*[ Let interviewee list wetlands, and double check for names not mentioned]*

| Estero | Visited | Research | Recreation (explain) | Other (explain) | Last visit (year/month) |
|---|---|---|---|---|---|
| San Jorge | | | | | |
| La Salina | | | | | |
| San Francisquito | | | | | |
| La Cinita | | | | | |
| Almejas | | | | | |
| La Pinta | | | | | |
| Morúa | | | | | |
| La Cholla | | | | | |
| Cerro Prieto | | | | | |
| Las Lisas | | | | | |
| San Judas | | | | | |
| Bahía Adair | | | | | |
| Other | | | | | |

14. Research activities. From those wetlands in which you carried out research activities, can you tell us the nature of those activities, please focus on unpublished works as we can easily complete the interview with published papers.

| Estero | Year | Project | Main findings, main results |
|---|---|---|---|
| San Jorge | | | |
| La Salina | | | |
| San Francisquito | | | |
| La Cinita | | | |
| Almejas | | | |
| La Pinta | | | |
| Morúa | | | |
| La Cholla | | | |
| Cerro Prieto | | | |
| Las Lisas | | | |
| San Judas | | | |
| Bahía Adair | | | |
| Other | | | |

Do you know of any projects/ research activities other researchers are carrying out in coastal wetlands in the northern Gulf between el Borrascoso and San Francisquito?
15. During your visits have you observed any change in the aspects of wetland function that I will mention?
Please state your answer in the categories of Increase/ No change/ Decrease/ +1/0/-1.
Can you mention when you first observed this change.

| Wetland function | 2000 | 1990 | 1980 | 1970 | 1960 | 1950 | Wetland | Species | Detailed description of change |
|---|---|---|---|---|---|---|---|---|---|
| Primary productivity | | | | | | | | | |
| Habitat and nursery for plant and animal species | | | | | | | | | |
| Maintenance of characteristic plant communities | | | | | | | | | |
| Biological diversity | | | | | | | | | |
| Water quality maintenance/nutrient retention | | | | | | | | | |
| Sediment retention | | | | | | | | | |
| Channel morphology | | | | | | | | | |
| Coastal morphology (mouth width, sedimentation) | | | | | | | | | |
| Wetland hydrology | | | | | | | | | |
| Food chain support | | | | | | | | | |

16. What are the main causes you have observed or assume are responsible for the changes in wetland community and function?

17. ¿In your judgment, what were the biggest threats to wetlands in each decade between 1950 y 2000?

| |
|---|
| a) 2000 |
| b) 1990 |
| c) 1980 |
| d) 1970 |
| e) 1960 |
| f) 1950 |

18. What are some rare or uncommon species you have seen in the wetlands?
19. Is there any particular story or thought about coastal wetlands you would like to share?

#### Text S6. Environmental history of coastal wetlands in the Northern Gulf of California

The current configuration of coastal wetlands in the Northern Gulf formed during the Holocene, when the sea level rise reached its highest point ~ 5000 years ago, and silt deposited by currents and tides over the ancient Pleistocene coast (Ortlieb 1991). Remains of former estuaries, in the form of salt pans or playas are found several kilometers inland; these former estuarine areas are surrounded by numerous shell middens that attest their previous connection to the sea (Foster *et al.* 2008). These wetlands have remained saltwater systems, although the Northern Gulf was wetter in the early Holocene due to increased winter rainfall, as evidenced by paleoclimatic studies; modern rainfall patterns were not fully established until after 9000 yr BP and generally wet conditions prevailed until about 6000 yr BP (Metcalfe *et al.* 2000). Coastal wetlands have progressively become hotter, drier and likely more saline.

Coastal wetlands in the Northern Gulf have been used through hundreds to several thousand years by humans. Flaked stone artifacts indicate probable Middle Archaic (5500 – 3500 BP) to Late Archaic (3500-2000 BP) use of the area (Foster *et al.* 2008). Artifacts from the Hohokam (from circa A.D. 400-1400+), Yuman (between A.D. 700-1500) and Trincheras cultures (between A.D. 1000 to 1300) are found in shell middens that surround the edges of the estuaries, dunes, and inland and can extend several kilometers (Foster *et al.* 2008, 2012). These groups exploited the variety of food sources found in the wetlands including oysters, conchs, clams, sea cucumbers, and crabs that became available during low tide; vertebrates such as fish, sea turtles, dolphins, sea lions and whales could also be consumed and trapped after entering the wetland at high tide (Foster 1975, Foster *et al.* 2012).

The indigenous groups encountered by the first European explorers in 1539 likely visited the coast but didn’t have permanent encampments because of the lack of freshwater (Mitchell & Foster 2000). These groups included the Hia C’ed O’odham and the Tohono O'odham; the latter traveled occasionally from Southern Arizona to collect salt in the salt flats surrounding Estero La Cholla and to conduct ceremonies as late as the early decades of the 20$^{th}$ century (Broyles *et al.* 2006). Following the Spanish explorations, including Father Eusebio Kino travels through Northwest Sonora between 1698-1706, indigenous presence in the area decreased as groups adopted farming and animal husbandry, succumbed to disease or were driven out (Broyles *et al.* 2006).

Modern occupation of the Northern Gulf began in the 1900’s, as seasonal fishing camps were established as year-round communities in Puerto Peñasco, Golfo de Santa Clara and San Felipe (Munro Palacio 2007). Demand for Totoaba (*Totoaba macdonaldi*) drove the start of commercial exploitation of fishery species in the Northern Gulf; as demand for Totoaba decreased fishers shifted to extracting shrimp (Cudney-Bueno &

Turk Boyer 1998). The industrial trawl shrimp fishery grew as the main fishery in the region until the 1980's, and this industry stimulated the growth of the small-scale fishing sector (Brusca & Bryner 2004). Today, 10,000-24,000 small-scale fishing boats operate daily in the Gulf and target over 70 species of fish and shellfish, having high ecological and economic impacts (Ezcurra *et al.* 2009). In the Northern Gulf, most fishing activities inside wetlands take place from *pangas*, small skiffs with outboard motors (*pangas*), using gill nets, traps, and cast nets (Cinti *et al.* 2010, Turk-Boyer *et al.* 2014); trawls are banned for use inside coastal wetlands (DOF 2013).

By the time the first aerial photographs of coastal wetlands were taken in 1956 (Dr. Joseph R. Schreiber, personal communication), anthropogenic modifications were evident in these coastal wetlands. Sometime between 1946, when the first description of Estero Morúa was published (Gifford 1946), and 1959, the course of the ephemeral Sonoyta river was diverted East due to the construction of a road, so that it no longer emptied into Estero Morúa (Foster 1975). In addition, starting in 1952 the Mexican government encouraged upstream groundwater pumping and diversion of the Sonoyta river for agriculture; such that the river no longer reaches the Gulf of California during the rainy season (Murguía-Ruiz 2001).

Other coastal wetlands impacted by tourism and development included Estero Puerto Peñasco which was dredged in 1972 and converted to a marina (Munro Palacio 2007). Development along the coast in Sonora intensified after 1993, when the Mexican constitution was modified to allow the division of *ejidos*, which owned most coastal lands, into individual parcels that could be bought and sold (Bracamonte-Sierra *et al.* 2008). For example, road area increased >400% within wetlands in the corridor between 1973-1997 (Sierra & Chamberlain 1999). In 1999, over 50% of Estero La Cholla was modified for the construction of a golf course and a dike as part of the Laguna del Mar residential development (Glenn *et al.* 2006). Starting in 2004, Estero La Pinta was extensively modified by dredging and filling during construction of a tourism resort (formerly the Mayan Palace Resort, now Grand Mayan - Vidanta) (personal observation). Based on available satellite images, by 2007 ~660 ha of this wetland had been filled or dredged, equivalent to 20% of the original wetland area (Morzaria-Luna *et al.* 2014). Trash, soil compaction, erosion and the construction of a coastal highway that transversed Bahía Adair are also problematic. The first descriptions of coastal wetlands in the Northern Gulf were published in 1994, with a detailed vegetation classification and analysis of change (1973-1997) (Cervantes 1994, Sierra & Chamberlain 1999). This analysis was updated (Brusca *et al.* 2006, Glenn *et al.* 2006) to include a photographic atlas (Nagler *et al.* 2004).

Currently, wetlands in Mexico are public domain and are administered as part of the Maritime-Terrestrial Federal Zone, which extends up to 20 meters inland from the highest tide line (Cortina-Segovia et al. 2007). Land tenure surrounding the wetlands in the Puerto Peñasco coastal corridor includes communal and private property; land use includes tourism, residential, fishing camps, and aquaculture (Morzaria-Luna et al. 2014a). Bahia Adair, Estero Cerro Prieto, and Estero La Cholla were included within the boundaries of the Upper Gulf of California and Colorado River Delta Biosphere Reserve, which was declared in 1993 (Conanp 2012) and named Ramsar Wetlands of International Importance in 2009. The coastal wetlands within Bahia San Jorge (Esteros La Salina, Almejas, and San Francisquito) were named Priority Terrestrial Regions for Biodiversity and Conservation (Arriaga et al. 2000) and Estero La Salina was included as part of Mexico's Important Bird Areas Program (AICA No. 34; (Conabio 2004). and Ramsar Wetlands of International Importance (Conanp 2012).

#### References

Bracamonte-Sierra, A., S.E. Meza-Martínez & R. Méndez-Barrón. 2008. Auge, crisis y perspectivas de Puerto Peñasco como destino turístico internacional. Topifilia. Revista de Arquitectura, Urbanismo y Ciencias Sociales Centro de Estudios de América del Norte, El Colegio de Sonora. 1.

Broyles, B., A.G. Rankin & R.S. Felger. 2006. Native peoples of the dry borders region, p. 128–146. *In* R.S. Felger & B. Broyles (eds.). . University of Utah Press, Salt Lake City, Utah.

Brusca, R.C. & G.C. Bryner. 2004. A case study of two Mexican biosphere reserves: The Upper Gulf of California and Colorado River Delta and the El Pinacate and Gran Desierto de Altar Biosphere Reserves, p. 28–64. *In* N.E. Harrison & G.C. Bryner (eds.). . Rowman & Littlefield Publishers, Lanham, MD.

Brusca, R.C., R. Cudney-Bueno & M. Moreno-Báez. 2006. Gulf of California esteros and estuaries. Analysis, state of knowledge, and conservation and priority recommendations. Arizona-Sonora Desert Museum.

Cervantes, M. 1994. Guía regional para el conocimiento, manejo y utilización de los humedales del Noroeste de México. CECARENA, Coordinación para la gestión de los humedales en México, Guaymas, Sonora.

Cinti, A., W. Shaw, R. Cudney-Bueno & M. Rojo. 2010. The unintended consequences of formal fisheries policies: Social disparities and resource overuse in a major fishing community in the Gulf of California, Mexico. Mar. Policy 34: 328–339.

Cudney-Bueno, R. & P.J. Turk Boyer. 1998. Pescando entre mareas del Alto Golfo de California: Una guía sobre pesca artesanal, su gente y sus propuestas de manejo. CEDO Intercultural, A.C., Puerto Peñasco, Sonora. México.

DOF. 2013. NORMA Oficial Mexicana NOM-002-SAG/PESC-2013, Para ordenar el aprovechamiento de las especies de camarón en aguas de jurisdicción federal de los Estados Unidos Mexicanos. Diario Oficial de la Federación. 11/07/2013.

Ezcurra, E., O. Aburto-Oropeza, M. de los Angeles Carvajal, R. Cudney-Bueno & J. Torre. 2009. Gulf of California, Mexico, p. 227–252. *In* K. McLeod & H. Leslie (eds.). Ecosystem-Based Management for the Oceans. Island Press.

Foster, J.W. 1975. Shell Middens, Paleoecology, and Prehistory: The Case from Estero Morua, Sonora, Mexico. Kiva 41: 185–194.

Foster, M.S., D.R. Mitchell, G. Huckleberry, D. Dettman & K.R. Adams. 2012. Archaic Period Shell Middens, Sea-Level Fluctuation, and Seasonality: Archaeology along the Northern Gulf of California Littoral, Sonora, Mexico. Am. Antiq. 77: 756–772.

Foster, M.S., D.R. Mitchell, G. Huckleberry & D.L. Dettman. 2008. Observations on the archaeology, paleoenvironment, and geomorphology of the Puerto Peñasco area of northern Sonora, Mexico. Kiva 73: 263–290.

Gifford, E.W. 1946. Archaeology in the Punta Penasco region, Sonora. Am. Antiq. 11: 215–221.

Glenn, E.P., P.L. Nagler, R.C. Brusca & O. Hinojosa-Huerta. 2006. Coastal wetlands of the northern Gulf of California: inventory and conservation status. Aquatic Conservation-Marine and Freshwater Ecosystems 16: 5–28.

Metcalfe, S.E., S.L. O'Hara, M. Caballero & S.J. Davies. 2000. Records of Late Pleistocene–Holocene climatic change in Mexico — a review. Quat. Sci. Rev. 19: 699–721.

Mitchell, D.R. & M.S. Foster. 2000. Hohokam Shell Middens along the Sea of Cortez, Puerto Penasco, Sonora, Mexico. J. Field Archaeol. 27: 27–41.

Morzaria-Luna, H.N., A. Castillo-López, G.D. Danemann & P.J. Turk-Boyer. 2014. Conservation strategies for coastal wetlands in the Gulf of California, Mexico. Wetlands Ecol. Manage. 22: 267–288.

Munro Palacio, G. 2007. Breve historia de Puerto Peñasco. De Cierto Mar Editores, Puerto Peñasco, Sonora.

Murguía-Ruiz, M.D.L. 2001. Water in El Pinacate y Gran Desierto de Altar Biosphere Reserve, Sonora, Mexico: Communities, wildlife and the border with the United States. Nat. Resour. J. 40: 411–434.

Nagler, P.L., E.P. Glenn & R.C. Brusca. 2004. Photographic atlas of the Gulf of California wetlands. University of Arizona.

Ortlieb, L. 1991. Quaternary Vertical Movements Along the Coasts of Baja California and Sonora: Chapter 22: Part III. Regional Geophysics and Geology, p. 447–480. *In* J.P. Dauphin & B.T. Simoneit (eds.). The Gulf and Peninsular Province of the Californias, Amer. Asçoc. Petrol. Geol. Mem. AAPG Special Volumes.

Sierra, R. & J. Chamberlain. 1999. Mapping and monitoring of coastal wetlands in Sonora, Mexico: a multi-national approach. Department of Geography. Arizona State University.

Turk-Boyer, P.J., H.N. Morzaria-Luna, I. Martinez-Tovar, C. Downton-Hoffmann & A. Munguia-Vega. 2014. Ecosystem-Based Fisheries Management of a Biological Corridor Along the Northern Sonora Coastline (NE Gulf of California), p. 125–154. *In* F. Amezcua & B. Bellgraph (eds.). Fisheries Management of Mexican and Central American Estuaries. Springer Netherlands, Dordrecht.

**Table S1. Fuzzy logic decision table.** Table summarizes the criteria used by the fuzzy logic algorithm to convert linguistic categories derived from fishers' interviews into relative abundance estimates for species (grouped as non-habitat dependent and habitat-dependent) and time periods. For each time period there are three low/high-yes/no ecosystem indicators (loss of wetland area, acknowledged depletion, and changes in breeding area). The interview abundance scores correspond to the abundance ratings derived from interviews, as low (L), medium-low(ML), medium (M), medium-high (MH), and high (H). The conclusions resulting from the combination of linguistic variables and indicators for each time period are indicated on a 0 (lowest) - 10 (highest) scale.

| Time period | Wetland loss | Depletion indicator | Breeding | Non-habitat dependent species | | | | | Habitat dependent species | | | | |
|---|---|---|---|---|---|---|---|---|---|---|---|---|---|
| | | | | Interview abundance score (linguistic category) | | | | | | | | | |
| | | | | L | ML | M | MH | H | L | ML | M | MH | H |
| 1950 | Low | NO | YES | 5 | 7 | 9 | 10 | 10 | 5 | 7 | 9 | 10 | 10 |
| | | YES | NO | 5 | 7 | 9 | 10 | 10 | 5 | 7 | 9 | 10 | 10 |
| | | | YES | 6 | 8 | 10 | 10 | 10 | 6 | 8 | 10 | 10 | 10 |
| | High | NO | NO | 2 | 4 | 6 | 8 | 10 | 6 | 8 | 10 | 10 | 10 |
| | | | YES | 5 | 7 | 9 | 10 | 10 | 7 | 9 | 10 | 10 | 10 |
| | | YES | NO | 5 | 7 | 9 | 10 | 10 | 7 | 9 | 10 | 10 | 10 |
| | | | YES | 6 | 8 | 10 | 10 | 10 | 9 | 10 | 10 | 10 | 10 |
| 1960 | Low | NO | YES | 4 | 6 | 8 | 10 | 10 | 4 | 6 | 8 | 10 | 10 |
| | | YES | NO | 4 | 6 | 8 | 10 | 10 | 4 | 6 | 8 | 10 | 10 |
| | | | YES | 5 | 7 | 9 | 10 | 10 | 5 | 7 | 9 | 10 | 10 |
| | High | NO | NO | 2 | 4 | 6 | 8 | 10 | 5 | 7 | 9 | 10 | 10 |
| | | | YES | 4 | 6 | 8 | 10 | 10 | 6 | 8 | 10 | 10 | 10 |
| | | YES | NO | 4 | 6 | 8 | 10 | 10 | 6 | 8 | 10 | 10 | 10 |
| | | | YES | 5 | 7 | 9 | 10 | 10 | 8 | 10 | 10 | 10 | 10 |
| 1970 | Low | NO | YES | 2 | 4 | 6 | 8 | 10 | 2 | 4 | 6 | 8 | 10 |
| | | YES | NO | 2 | 4 | 6 | 8 | 10 | 2 | 4 | 6 | 8 | 10 |
| | | | YES | 1 | 3 | 5 | 7 | 9 | 1 | 3 | 5 | 7 | 9 |
| | High | NO | NO | 2 | 4 | 6 | 8 | 10 | 2 | 4 | 6 | 8 | 10 |
| | | | YES | 2 | 4 | 6 | 8 | 10 | 1 | 3 | 5 | 7 | 9 |
| | | YES | NO | 2 | 4 | 6 | 8 | 10 | 1 | 3 | 5 | 7 | 9 |
| | | | YES | 1 | 3 | 5 | 7 | 9 | 0 | 2 | 4 | 6 | 8 |
| 1980 | Low | NO | YES | 0 | 2 | 4 | 6 | 8 | 0 | 2 | 4 | 6 | 8 |
| | | YES | NO | 0 | 2 | 4 | 6 | 8 | 0 | 2 | 4 | 6 | 8 |
| | | | YES | 0 | 1 | 3 | 5 | 7 | 0 | 1 | 3 | 5 | 7 |
| | High | NO | NO | 2 | 4 | 6 | 8 | 10 | 0 | 2 | 4 | 6 | 8 |
| | | | YES | 0 | 2 | 4 | 6 | 8 | 0 | 1 | 3 | 5 | 7 |

| | | | | | | | | | | | | | |
|---|---|---|---|---|---|---|---|---|---|---|---|---|---|
| | | YES | NO | 0 | 2 | 4 | 6 | 8 | 0 | 1 | 3 | 5 | 7 |
| | | | YES | 0 | 1 | 3 | 5 | 7 | 0 | 0 | 2 | 4 | 6 |
| 1990 | Low | NO | YES | 0 | 1 | 3 | 5 | 7 | 0 | 1 | 3 | 5 | 7 |
| | | YES | NO | 0 | 1 | 3 | 5 | 7 | 0 | 1 | 3 | 5 | 7 |
| | | | YES | 0 | 0 | 2 | 4 | 6 | 0 | 0 | 2 | 4 | 6 |
| | High | NO | NO | 2 | 4 | 6 | 8 | 10 | 0 | 1 | 3 | 5 | 7 |
| | | | YES | 0 | 1 | 3 | 5 | 7 | 0 | 0 | 2 | 4 | 6 |
| | | YES | NO | 0 | 1 | 3 | 5 | 7 | 0 | 0 | 2 | 4 | 6 |
| | | | YES | 0 | 0 | 2 | 4 | 6 | 0 | 0 | 1 | 3 | 5 |
| 2000 | Low | NO | YES | 0 | 0 | 2 | 4 | 6 | 0 | 0 | 2 | 4 | 6 |
| | | YES | NO | 0 | 0 | 2 | 4 | 6 | 0 | 0 | 2 | 4 | 6 |
| | | | YES | 0 | 0 | 1 | 3 | 5 | 0 | 0 | 1 | 3 | 5 |
| | High | NO | NO | 2 | 4 | 6 | 8 | 10 | 0 | 0 | 2 | 4 | 6 |
| | | | YES | 0 | 0 | 2 | 4 | 6 | 0 | 0 | 1 | 3 | 5 |
| | | YES | NO | 0 | 0 | 2 | 4 | 6 | 0 | 0 | 1 | 3 | 5 |
| | | | YES | 0 | 0 | 1 | 3 | 5 | 0 | 0 | 0 | 2 | 4 |
| All decades | Low | NO | NO | 2 | 4 | 6 | 8 | 10 | 2 | 4 | 6 | 8 | 10 |

**Table S2. List of bibliographic references describing scientific studies carried out within wetlands in the Northern Gulf or that used samples collected inside them.**

| Citation |
|---|
| 1. Alatorre-Sanchez J. R. 2008. Valoración Contingente Del Hábitat de Invierno de Las Aves Playeras Migratorias En La Costa Del Pacífico En América Del Norte. Instituto Tecnológico Autónomo de México, México, Distrito Federal. México. Tesis de Licenciatura en Economía |
| 2. Alison Leeds D. 1981. Analysis of Flow to Pumping Wells in a Saline Coastal Aquifer, Puerto Peñasco, Sonora, Mexico. The University of Arizona. Master's Thesis |
| 3. Anderson D. W., Henny C. J., Godinez-Reyes C., Palacios E. L., Santos del Prado K., Bredy J. 2007. Size of the California Brown Pelican Metapopulation during a Non-El Niño. Reston, Virginia: U.S. Geological Survey, Open-File Report 2007-1299. |
| 4. Ayala F., O' Leary J. W. 1995. Growth and Physiology of *Salicornia Bigelovii* Torr. at Suboptimal Salinity. *International Journal of Plant Sciences* 156 (2): 197–205. |
| 5. Ayala F., W. O'Leary J., Schumaker K. S. 1996. Increased Vacuolar and Plasma Membrane H+-ATPase Activities in *Salicornia Bigelovii* Torr. in Response to NaCl. *Journal of Experimental Botany* 47 (1): 25–32. |
| 6. Beu A. G., Knudsen J. 1986. Taxonomy of Gastropods of the Families Ranellidae (=Cymatiidae) and Bursidae. Part 3. A Review of the Trifid-Ribbed Species of Cymatium (Turritriton). *Journal of the Royal Society of New Zealand* 17 (1): 73–91. |
| 7. Boyer K. L., Barker C. L. 1971. An Investigation of the Physiological Ecology of the Gobiid Fish Quietula Guaymasiae in the Gulf of California. In , VIII:22pp ( from 22–42 pp). Biological Studies in the Gulf of California. University of Arizona. |
| 8. Brown J. J., Glenn E. P. 1999. Reuse of Highly Saline Aquaculture Effluent to Irrigate a Potential Forage Halophyte, Suaeda Esteroa. *Aquacultural Engineering* 20 (2): 91–111. |
| 9. Brown J. J., Glenn E. P., Fitzsimmons K. M., Smith S. E. 1999. Halophytes for the Treatment of Saline Aquaculture Effluent. *Aquaculture* 175 (3-4): 255–68. |
| 10. Bryant E. H. 2007. Barriers to Sustainable Coastal Development in Puerto Peñasco, Sonora, Mexico. Duke University.Master's project, Duke University. https://hdl.handle.net/10161/277. |

11. Bucciarelli G., Di Filippo M., Costagliola D., Alvarez-Valin F., Bernardi G., G. Bernardi. 2009. Environmental Genomics: A Tale of Two Fishes. *Molecular Biology and Evolution* 26 (6): 1235–43.

12. Cadee G. C., Walker S. E., Flessa K. W.. 1997. Gastropod Shell Repair in the Intertidal of Bahia La Choya (N. Gulf of California). *Palaeogeography, Palaeoclimatology, Palaeoecology* 136 (1-4): 67–78.

13. Carmona R., Hernández-Alvarez A., Martínez-Reséndiz B., Ruiz-Campos G., de la Cruz-Agüero J., Saldierna R, Cota-Gómez V.M., Hernández-Rivas M., Danemann G.D. 2017. Biología y Conservación Del Pejerrey (Atherinopsidae, *Leuresthes sardina*). *Ciencia Pesquera* 25 (2): 51–67.

14. Carmona-Islas C., Bello-Pineda J., Carmona R., Velarde E.. 2013. Modelo Espacial Para La Detección de Sitios Potenciales Para La Alimentación de Aves Playeras Migratorias En El Noroeste de México. *Huitzil* 14 (1): 22–36.

15. Cervantes M. 1994. Guía Regional Para El Conocimiento, Manejo Y Utilización de Los Humedales Del Noroeste de México. Guaymas, Sonora: CECARENA, Coordinación para la gestión de los humedales en México. 155 p.

16. Cheney D. P. 1992. Hydrology and Geochemistry of Modern, Supratidal Evaporite Deposits, Bahia Adair, Sonora, Mexico. Southwestern Louisiana University PhD Thesis.

17. Crawford C. S., Campbell M. L., Schaedla W. H., Wood S. 1989. Assemblage Organization of Surface-Active Arthropods along Horizontal Moisture Gradients in a Coastal Sonoran Desert Ecosystem. *Acta Zoologica Mexicana* 34: 30–51.

18. Cruz, V. M., del Río A.C., Ortlieb L. 1978. Transgresiones Cuaternarias En La Costa de Sonora. Universidad Nacional Autónoma de México. Inst. Geología *Revista Mexicana de Ciencias*. Vol. 2, 90-97.

19. Cutler A. H. 1987. Shell Microtextures as Records of Taphonomic History. *Geological Society of America. Programs with Abstracts* 19 (7): 634.

20. Cutler A. H. 1995. Taphonomic Implications of Shell Surface Textures in Bahia-La-Choya, Northern Gulf of California. *Palaeogeography, Palaeoclimatology, Palaeoecology* 114 (2-4): 219–40.

21. Dawson E. Y. 1966. Benthic Algae in the Northernmost Gulf of California, Mexico. In. Unpaginated).Academy of Sciences 4: 55–66

22. Dawson E. Y. 1966. Marine Algae in the Vicinity of Puerto Penãsco, Sonora, Mexico. Gulf of California Field Guide Series, No. 1. Tucson: University of Arizona.

23. Dawson E. Y. 1966. New Records of Marine Algae from the Gulf of California. *Journal of the Arizona-Nevada Academy of Science* 4 (2): 55–56.

24. Delgado-Estrella A., Ortega-Ortiz J. G., Sánchez-Ríos A. 1994. Varamiento de Mamíferos Marinos Durante Primavera Y Otoño, Y Su Relación Con La Actividad Humana En El Norte Del Golfo de California. *Anales Del Instituto de Biología, Universidad Nacional Autónoma de México, Serie Zoología* 65 (2): 287–95.

25. Dettman D. L., Mitchell D. R., Huckleberry G., Foster M.S. 2015. 14C and Marine Reservoir Effect in Archaeological Samples from the Northeast Gulf of California. *Radiocarbon* 57 (5): 785–93.

26. Donath-Hernandez F. E. 1988. Three New Species of Cumacea from the Gulf of California (Crustacea, Peracarida). *Cahiers de Biologie Marine*., 29 (4), 531–543.

27. Ekdale A. A. 1987. Late Cenozoic Rocks in the Puerto Penasco Area. *The Paleontological Society Special Publications* 2: 34–43.

28. Enríquez-Espinoza T. L., Grijalva-Chon J.M., Castro-Longoria R., Ramos-Paredes J. 2010. Perkinsus Marinus in *Crassostrea Gigas* in the Gulf of California. *Diseases of Aquatic Organisms* 89 (3): 269–73.

29. Ezcurra E., Felger R. S., Russell A. D., Equihua M. 1988. Freshwater Islands in a Desert Sand Sea: The Hydrology, Flora, and Phytogeography of the Gran Desierto Oases of Northwestern Mexico. *Desert Plants* 9 (2): 35–44, 55–63.

30. Felger R. 1992. Synopsis of the Vascular Plants of Northwestern Sonora, Mexico. *Ecologica* 2 (2): 11–44.

31. Felger R. S. 1980. Vegetation and Flora of the Gran Desierto, Sonora, México. Desert Plants 2(2): 0734-3434.

32. Felger R. S., Lowe C. H. 1976. The Island and Coastal Vegetation and Flora of the Northern Part of the Gulf of California. Contributions in Science; No 285. Los Angeles: Natural History Museum of Los Angeles County.

33. Flessa K. W., Cutler A. H, Meldahl K. H. 1993. Time and Taphonomy: Quantitative Estimates of Time-Averaging and Stratigraphic Disorder in a Shallow Marine Habitat. *Paleobiology* 19 (2): 266–86.

34. Flessa K. W., Ekdale A. A., Flessa K. W. 1987. Paleoecology and Taphonomy of Recent to Pleistocene Intertidal Deposits, Gulf of California. In , 2–33. The Paleontological Society Special Publication. Washington, D.C: Paleontological Society.

35. Foster J. W. 1975. Shell Middens, Paleoecology, and Prehistory: The Case

from Estero Morua, Sonora, Mexico. *The Kiva* 41 (2): 185–94.

36. Foster M. S., Mitchell D. R., Huckleberry G., Dettman D. L. 2008. Observations on the Archaeology, Paleoenvironment, and Geomorphology of the Puerto Peñasco Area of Northern Sonora, Mexico. *The Kiva* 73 (3): 263–90.

37. Foster M.S., Mitchell D.R., Huckleberry G., Dettman D., Adams K.R. 2012. Archaic Period Shell Middens, Sea-Level Fluctuation, and Seasonality: Archaeology along the Northern Gulf of California Littoral, Sonora, Mexico. *American Antiquity* 77 (4): 756–72.

38. Frias-Martins A. M. 1996.The anatomy of Cassidula Ferussac, 1821 and a case for the revival of the Cassidulinae Odhner, 1925. In: Morten B, editor. Proceedings of the Third International Conference on the Marine Biology of the South China Sea, The Marine Biology of the South China Sea. Hong Kong University Press, Hong Kong. 25-42.

39. Furisch F. T., Flessa K. W. 1986. Taphonomy of an Intertidal Mollusca Fauna; Implications for Palaeoecological Studies. 11. Conference Paper.

40. Fursich F. T., Flessa K., Aberhan M., Feige A., Schodelbauer S. 1991. Sedimentary Habitats and Molluscan Fauna of Bahia La Choya (Gulf of California, Sonora, Mexico). *Zitteliana* 18: 5–51.

41. Fürsich F. T., Flessa K.W. 1987. Taphonomy of Tidal Flat Molluscs in the Northern Gulf of California: Paleoenvironmental Analysis despite the Perils of Preservation. *The Paleontological Society Special Publications* 2 (6): 200–237.

42. Gallardo-Ybarra C., Minjarez-Osorio C., Grijalva-Chon J.M., Castro-Longoria R., Lastra-Encinas M.A., De-la-Re-Vega E. 2019. Expresión y tropismo del Herpesvirus de Ostreidos tipo 1 en dos tejidos del ostión del Pacífico *Crassostrea gigas*. *Biotecnia* 21 (3): 35–40.

43. Garcia-Rico L., Valenzuela Rodriguez M., Jara-Marini M.E. 2006. Geochemistry of Mercury in Sediment of Oyster Areas in Sonora, Mexico. *Marine Pollution Bulletin* 52 (4): 453–58.

44. García-Silva G., Marinone S.G. 1997. Modelado de Corrientes Residuales En El Golfo de California Mediante La Utilización de Diferentes Tamaños de Malla. *Ciencias Marinas* 23 (4): 505–19.

45. García-Hernández J., Hinojosa-Huerta O., Gerhart V., Carrillo-Guerrero Y., Glenn E. P. 2001. Willow Flycatcher (*Empidonax Traillii*) Surveys in the Colorado River Delta: Implications for Management. *Journal of Arid Environments* 49 (1): 161–69.

46. Glenn E. P., O'Leary W. 1984. Relationship between Salt Accumulation and

Water Content of Dicotyledonous Halophytes. *Plant, Cell & Environment* 7 (4): 253–61.

47. Glenn, E. P., O'leary J., Watson M.C., Thompson T.L., Kuehl R.O. 1991. *Salicornia Bigelovii* Torr.: An Oilseed Halophyte for Seawater Irrigation. *Science* 251 (4997): 1065–67.

48. Gomez-Sapiens M. M., Soto-Montoya E., Hinojosa-Huerta O. 2013. Shorebird Abundance and Species Diversity in Natural Intertidal and Non-Tidal Anthropogenic Wetlands of the Colorado River Delta, Mexico. *Ecological Engineering* 59 (October): 74–83.

49. González R. 1991. Archaeology as a Geologic Tool : Interpretation of the Holocene Sedimentary History of Bahia Adair, Sonora, Mexico. University of Louisiana. PhD Thesis.

50. González-Escobar M., Pérez-Tinajero C.I., Suárez-Vidal F., González-Fernández A. 2013. Structural Characteristics of the Altar Basin, Northwest Sonora, Mexico. *International Geology Review* 55 (3): 322–36.

51. Goodfriend W. L. 1998. Microbial Community Patterns of Potential Substrate Utilization: A Comparison of Salt Marsh, Sand Dune, and Seawater-Irrigated Agronomic Systems. *Soil Biology & Biochemistry* 30 (8-9): 1169–76.

52. Goodfriend W. L., Olsen M. W., Frye R. J. 2000. Soil Microfloral and Microfaunal Response to *Salicornia Bigelovii* Planting Density and Soil Residue Amendment. *Plant and Soil* 223 (1-2): 23–32.

53. Goodfriend W. L., Olsen M. W., Frye. 1998. Decomposition of Seawater-Irrigated Halophytes - Implications for Potential Carbon Storage. *Plant & Soil* 202 (2): 241–50.

54. Grismer L. L. 1994. Three New Species of Intertidal Side-Blotched Lizards (*Genus Uta*) from the Gulf of California, México. *Herpetologica* 50 (4): 451–74.

55. Hall J. D. 1966. The Effect of the Venom of the Round Stingray Urobatis Halleri Found in Cholla Bay. In , IV:12. Biological Studies in the Gulf of California.Vol. IV, No. 2 (collected student papers from U. of Ariz. Marine Ecology class), 12 pp. @ UCSD(S/R)-UA(S)

56. Hamilton, D. C. 1996. Interpretation of Groundwater Salinity in a Coastal, Evaporitic Environment, Bahia Adair, Sonora, Mexico.

57. Hayes M. J., Flessa K. W. 1987. Surficial Physical Sedimentary Structures of Bahia La Choya. In , 52–61. The Paleontological Society Special Publication. Washington, D.C: Paleontological Society.

58. Hendrickx M. E. 1995. Checklist of Lobster-like Decapod Crustaceans (Crustacea:Decapoda:Thalassinidea. Astacidea and Palinuridea) from the Eastern Tropical Pacific. *Anales Del Instituto de Biología, Universidad Nacional Autónoma de México, Serie Zoología* 66 (002): 151–63.

59. Hendry D., Ekdale A.A. 1987. Color Pattern Variation in Populations of T*heodoxus Luteofasciatus* in the Puerto Penasco Area. *The Paleontological Society Special Publications* 2: 104–12.

60. Herbert G. S. 2004. Observations on Diet and Mode of Predation in *Stramonita Biseriali*s (*Gastropoda: Muricidae*) from the Northern Gulf of California. *Festivus*. Vol.36. 41-45.

61. Hiriart J.L. 1989. Crustáceos estomatópodos y decápodos intermareales de las islas del Golfo de California, México. UNAM.

62. Hollenberg G. J., Norris J. N. 1977. The Red Alga Polysiphonia (Rhodomelaceae) in the Northern Gulf of California. Smithsonian Contributions to the Marine Sciences. Smithsonian Institution.

63. Huang D., Bernardi G. 2001. Disjunct Sea of Cortez-Pacific Ocean *Gillichthys Mirabilis* Populations and the Evolutionary Origin of Their Sea of Cortez Endemic Relative, *Gillichthys Seta*. *Marine Biology* 138 (2): 421–28.

64. Huey L. M. 1935. February Bird Life of Punta Penascosa, Sonora, Mexico. *The Auk* 52 (3): 249–56.

65. Johnson A. F. 1982. Dune Vegetation along the Eastern Shore of the Gulf of California. *Journal of Biogeography* 9 (4): 317–30.

66. Kearney W. S., Fagherazzi S. 2016. Salt Marsh Vegetation Promotes Efficient Tidal Channel Networks. *Nature Communications* 7: 12287.

67. Kudenov, Jerry D. 1975. Errant Polychaetes from the Gulf of California, Mexico. *Journal of Natural History* 9 (1): 65–91.

68. Lamb R. R. 1970. A Survey of the Blue Crab, *Callinectes Bellicosus*, from the Estero Morua, Sonora, Mexico. In , VII:10. Biological Studies in the Gulf of California. No. 2556.

69. Lancaster N. 1992. Relations between Dune Generations in the Gran Desierto of Mexico. *Sedimentology* 39 (4): 631–44.

70. Lane J. D. 1964. Descriptions of and a Key to the Gobiid Fishes Found near Puerto Penasco, Sonora, Mexico. I:7. Biological Studies in the Gulf of California.

71. Lau C. L., Jacobs D.K. 2017. Introgression between Ecologically Distinct Species Following Increased Salinity in the Colorado Delta- Worldwide Implications for Impacted Estuary Diversity. *PeerJ* 5 (December): e4056.

72. Lewis D. E. Jr. 1977. Standing Crop Biomass of Intertidal Plants in Estero de La Cholla, Sonora, Mexico. In , XII:24. Biological Studies in the Gulf of California.

73. León-González J. A. 1998. Informe Final Del Proyecto H011 Nereididae (Annelida: Polychaeta) de México. México, D.F.: Universidad Autónoma de Nuevo León, Facultad de Ciencias Biológicas Departamento de Zoología de Invertebrados, Laboratorio de Zoología de Invertebrados No-Arthropoda.

74. Lock B. E. 2002. Sabkhas Ancient and Modern. Gulf Coast Association of Geological Societies Transactions. 52:645-657

75. Lock, B. E., Sinitiere S.M., Williams L.J. 1989. Bahia Adair and Vicinity, Sonora: Modern Siliciclastic-Dominated Arid Macrotidal Coastline. 73:3. Univ. of Southwestern Louisiana, Lafayette (USA).

76. Long G. E., Taylor W. 1969. Estuarine Mollusks of the Cholla Bay, Sonora, Mexico. In *The Echo: Abstracts and Proceedings of the Second Annual Meeting of the Western Society of Malacologists*, 2:17–18.

77. Luévano-Esparza J., Delgadillo-Vásquez A.M., Montes-Ontiveros O. 2015. Estructuras Artificiales Para La Anidación Y Su Relación Con El éxito Reproductivo Del Gavilán Pescador Y Del Tecolote Llanero Durante Ocho Temporadas Reproductivas En El Estero La Pinta, Puerto Peñasco, Sonora, México. *Huitzil* 16 (1): 9–15.

78. Mabry, J. B., Brusca R.C., Brack M.L., Miljour H.J., Gaines E.P. 2007. Preliminary Report for the Project ``Fechamiento de Concheros Prehistóricos Del Estero Morua, Sonora, México''. rickbrusca.com.

79. Mackie S., Mackie M.. 1988. Cangrejos Violinistas En Estero Morua. (Fiddler Crabs in Estero Morua). *CEDO News* 1 (2): 14–17.

80. Magar V., González-García L., Markus S. Gross. 2017. Evaluación técnico-económica del potencial de desarrollo de parques eólicos en mar: el caso del Golfo de California. *Biotecnia* 19 (0): 3–8.

81. Mathews, C., and J. Balmer. 1971. Habitat and Behavior of Three Species of Fiddler Crabs Encountered in Puerto Penasco and Guaymas, Mexico. In , VIII:8. Biological Studies in the Gulf of California.

82. May L. A. 1973. Geological Reconnaissance of the Gran Desierto Region, Northwestern Sonora, Mexico. *Journal of the Arizona-Nevada Academy of Science* 8 (3): 158–69.

83. Mc Court R. M. 1984. Niche differences between sympatric Sargassum species in the northern Gulf of California. Marine Ecology Progress Series 18: 139-148.

84. Meldahl K. H. 1987. Biogenic and Physical Modes of Stratification and Shell Bed Formation, Bahia La Cholla, Northern Gulf of California. *Geological Society of America. Programs with Abstracts* 19 (7): 770.

85. Meldahl Keith H. 1987. Sedimentologic and Taphonomic Implications of Biogenic Stratification. *Palaios* 2 (4): 350–58.

86. Mitchell D. R., Foster M. S. 2000. Hohokam Shell Middens along the Sea of Cortez, Puerto Penasco, Sonora, Mexico. *Journal of Field Archaeology* 27 (1): 27–41.

87. Mitchell D. R., Huckleberry G., Rowell K., Dettman D.L. 2015. Coastal Adaptations During the Archaic Period in the Northern Sea of Cortez, Mexico. *The Journal of Island and Coastal Archaeology* 10 (1): 28–51.

88. Molina Ocampo R. E., Cisneros Mata M. A., Belindez Moreno L. F., Zarate Becerra E., Gaspar Dillanes M. T., Lopez Gonzalez L. C., Saucedo Ruiz C., Tovar Ávila J. 1999. La Jaiba En Sonora. In , 327–48. Ensenada, BC: Instituto Nacional de La Pesca. Centro Regional de Investigación Pesquera.

89. Morzaria-Luna H. N., Castillo-López A., Danemann G. D., Turk-Boyer P. J.. 2014. Conservation Strategies for Coastal Wetlands in the Gulf of California, Mexico. *Wetlands Ecology and Management* 22 (3): 267–88.

90. Morzaria-Luna H. N., Cruz-Piñón G., Brusca R. C., López-Ortiz A. M., Moreno-Báez M., Reyes-Bonilla H., Turk-Boyer P. J. 2018. Biodiversity Hotspots Are Not Congruent with Conservation Areas in the Gulf of California. *Biodiversity and Conservation* 27 (14): 3819–42.

91. Morzaria-Luna H., Turk-Boyer P. J., Rosemartin A., Camacho-Ibar V. F. 2014. Vulnerability to Climate Change of Hypersaline Salt Marshes in the Northern Gulf of California. *Ocean & Coastal Management* 93: 37–50.

92. Morzaria-Luna, H., Iris-Maldonado A., Valdivia-Jiménez P. 2010. Physico-chemical characteristics of negative estuaries in the northern Gulf of California, Mexico. in, J. R. Crane and A. E . Solomon (eds), Esutuaries: Types, Movement Patterns and Critical Impacts. Nova Science Publishers, 201-223 pp.

93. Munro G. 1985. La Leyenda de Puerto Peñasco. *Técnica Pesquera*, 6–10.

94. Norris J. N., Johansen H. W. 1981. Articulated Coralline Algae of the Gulf of California, Mexico, I: *Amphiroa* Lamouroux. Washington: Smithsonian Institution Press.

95. Ortlieb L. 1991. Quaternary Vertical Movements Along the Coasts of Baja California and Sonora: Chapter 22: Part III. Regional Geophysics and Geology. In *The Gulf and Peninsular Province of the Californias*, edited by J. P. Dauphin and B. T. Simoneit, 447–80. *Amer. Asçoc. Petrol. Geol. Mem.*

AAPG Special Volumes.

96. Owen W., Owens M. W. 1972. An Analysis of Digging Sites of *Callinectes Bellicosus* in Cholla Bay, Sonora, Mexico. IX:4. Biological Studies in the Gulf of California. Collected Student Papers. University of Arizona. *Mar. Ecol*. IX: 4 pp.

97. Pagel C. 1970. Planktonic population of Estero Morua and Station Beach: A comparative study. Biological Studies in the Gulf of California, Vol. VII, No. 2 (collected student papers from U. of Ariz. Marine Ecology Class), 5 pp. @ UCSD(S/R)-UA(S)

98. Palacios E., Mellink E. 1996. Status of the Least Tern in the Gulf of California (Estado de Sterna Antillarum En El Golfo de California). *Journal of Field Ornithology* 67 (1): 48–58.

99. Place S. P., Hofmann G.E. 2001. Temperature Interactions of the Molecular Chaperone Hsc70 from the Eurythermal Marine Goby *Gillichthys mirabilis*. *The Journal of Experimental Biology* 204 (Pt 15): 2675–82.

100. Puente E.O., Beltrán F., Ruiz F., Valdez R., García J., Avila N., Partida L., Murillo B. 2010. Sustainable Options for Soil Management in Arid Zones: Uses of the Halophyte *Salicornia Bigelovii* (torr.) and Biofertilizers in Modern Agriculture. *Tropical and Subtropical Agroecosystems* 13 (2): 157–67.

101. Rico L.G., Soto Cruz M.S., Jara Marini M.E., Gómez Álvarez A. 2004. Fracciones Geoquímicas de Cd, Cu Y Pb En Sedimentos Costeros Superficiales de Zonas Ostrícolas Del Estado de Sonora, México. *Revista Internacional de Contaminación Ambiental* 20 (4): 159–67.

102. Rodríguez-Pérez M., Aurioles-Gamboa D., Sánchez-Velásco L., Lavín M.F., Newsome S.D.. 2018. Identifying Critical Habitat of the Endangered Vaquita (*Phocoena Sinus*) with Regional δ 13 C and δ 15 N Isoscapes of the Upper Gulf of California, Mexico : The critical habitat of vaquita. *Marine Mammal Science* 34 (3): 790–805.

103. Rosales-Hoz L, Carranza-Edwards A., Aguirre-Gomez A., Galán-Alcala A. 1988. Estudio de Metales En Sedimentos Litorales de Sonora, México. *Anales Del Instituto de Ciencias Del Mar y Limnología, Universidad Nacional Autónoma de México* 15 (2): 225–34.

104. Rose M. 1976. Sedimentology of Estero La Cholla, Northwest Coast of Sonora, Mexico. University of Arizona. Master´s Thesis.

105. Rueda-Puente E. O., Villegas-Espinoza J.A., Gerlach-Barrera L.E., Tarazón-Herrera M.A., Murillo-Amador B., García-Hernández J.L., Troyo-Diéguez E., Preciado-Rangel P. 2009. Efecto de La Inoculación de Bacterias Promotoras de Crecimiento Vegetal Sobre La Germinación de *Salicornia bigelovii*. *Terra Latinoam* 27 (4): 345–54.

106. Sandusky C. 1969. Sedimentology of Estero Marua, Sonora, Mexico. Univ. of Arizona, M.S. Thesis

107. Scheidt S., Lancaster N., Ramsey M. 2011. Eolian Dynamics and Sediment Mixing in the Gran Desierto, Mexico, Determined from Thermal Infrared Spectroscopy and Remote-Sensing Data. *Geological Society of America Bulletin* 123 (7-8): 1628–44.

108. Schmidt N. 1988. An Examination of Shell Repair in the Northern Gulf of California. *Western Society of Malacology Annual Report (for 1987)* 20: 26.

109. Schultz G. A. [1]. 1970. A Review of the Species of the Genus Tylos Latreille From the New World (*Isopoda, Oniscoidea*). *Crustaceana* 19: 297–305.

110. Segura E. P., Jacques-Ayala C., César. 1991. *Studies of Sonoran Geology*. Geological Society of America. Geological Society of America. Vol. 254. DOI: https://doi.org/10.1130/SPE254.

111. Siegel-Causey D. 1981. Specific Affinities of the Gammaridean Fauna of Estero Morúa, Sonora, Mexico. In , XV:25. Biological Studies in the Gulf of California.

112. Sinitiere S. M. 1989. The Origin of a Late Quaternary Non-Marine Evaporite Sequence in the Gran Desierto/Bahia Adair Region, Sonora, Mexico. M.Sc. Thesis, University of Southwestern Louisiana

113. Skoglund C. C. 1965. Gastropods of Cholla Bay, Sonora, Mexico. In , II:10. Biological Studies in the Gulf of California. Vol, 11 (collected student papers from U. of Ariz. Marine Ecology Class), 17 pp. @ UCSD(S/R)-UA(S)

114. Skoglund C. C. 1974. Distribution and Associations of Eight Species of Intertidal Nassarius at Cholla Bay, Sonora, Mexico. *ECHO*, no. 6: 27.

115. Skoglund C. C. 1977. Egg Capsules of Five Species of Nassarius (Gastropoda) from Bahia Cholla, Sonora, Mexico. *Western Society of Malacology Annual Report* 10: 13–15.

116. Springer D. A., Flessa K. W. 1996. Faunal Gradients in Surface and Subsurface Shelly Accumulations from a Recent Clastic Tidal Flat, Bahia La Choya, Northern Gulf of California, Mexico. *Palaeogeography, Palaeoclimatology, Palaeoecology* 126 (3-4): 261–79.

117. Straw J. L. 1967. Movement of Fiddler Crabs at Puerto Penasco. In , V:10. Biological Studies in the Gulf of California. Vol. V, No. 1 (collected student papers from U. of Ariz. Marine Ecology Class), 10 pp. @ UCSD(S/R)-UA(S)

118. Sumpter L. T., Flessa K. W. 1987. Grain Size and Provenance of Bahia

La Choya Sediments. In , 44–51. The Paleontological Society Special Publication. Washington, D.C: Paleontological Society.

119. Swift C. C., Findley Ll. T., Ellingson R.A., Flessa K.W., Jacobs D.K. 2011. The Delta Mudsucker, *Gillichthys Detrusus*, a Valid Species (Teleostei: Gobiidae) Endemic to the Colorado River Delta, Northernmost Gulf of California, Mexico. *Copeia* 2011 (1): 93–102.

120. Swingle R. S., Glenn E. P., Squires V. 1996. Growth Performance of Lambs Fed Mixed Diets Containing Halophyte Ingredients. *Animal Feed Science and Technology* 63 (1-4): 137–48.

121. Titley J. 1982. Evidence That Housing May Be a Limiting Factor for Population Densities of *Octopus Digueti* in Cholla Bay, Sonora, Mexico. in: XVI:8. Biological Studies in the Gulf of California.

122. Turk-Boyer P. J., Morzaria-Luna H.N., Martinez-Tovar I., Downton-Hoffmann C., Munguia-Vega A. 2014. Ecosystem-Based Fisheries Management of a Biological Corridor Along the Northern Sonora Coastline (NE Gulf of California). In *Fisheries Management of Mexican and Central American Estuaries*, edited by Amezcua F., Bellgraph B., 125–54. Dordrecht: Springer Netherlands.

123. Turk-Boyer P.J., Peña-Bonilla H., Morzaria-Luna H.N., Valdivia-Jiménez P.A., Tovar-Vázquez H., Castillo-López A. 2014. Wetland Conservation in Northern Sonora, Mexico: Legal Tools and Active Communities. In *Fisheries Management of Mexican and Central American Estuaries*, edited by Felipe Amezcua and Brian Bellgraph, 183–206. Estuaries of the World. Dordrecht: Springer Netherlands.

124. Valdes-Casillas C., Carrillo-Guerrero Y., Zamora-Arroyo F., Hinojosa-Huerta O., Camacho-López M., Delgado-García S., Moreno-Báez M. 1999. Mapping and Management of Coastal Wetlands of Puerto Peñasco, Sonora: A Multinacional Project. Sonora: Center for Conservation of Natural Resources (CECARENA), Instituto Tecnológico y de Estudios Superiores de Monterrey -- Campus Guaymas (ITESM-CG); Pronatura. Arizona State University.

125. Verdugo-Fimbres M. I. 1985. *Presente Y Pasado: Historia Del Municipio de Puerto Peñasco*. Hermosillo, Sonora. INAH-SEP. Centro Regional del Noroeste, 1985 - 86 páginas

126. Villalobos-Hiriart J. L., Nates-Rodriguez J. C., Diaz-Barriaga A. C., Valle-Martinez M. A., Hernandez-Flores P., Lira-Fernandez E., Schmidtsdorf-Valencia P. 1989. Listados Faunísticos de México. I Custaceos Estomatopodos y Decápodos Intermareales de Las Islas Del Golfo de California, México. México, D.F.: Universidad Nacional Autónoma de México Instituto de Biología, UNAM.

127. Voight J. R. 1984. Habitat Selection, Reproduction and Sibling Species of *Octopus Digueti*. *Journal of the Arizona-Nevada Academy of Science* 19: 13–14.

128. Voight J. R. 1986. Octopus Research in the Gulf of California. *Newsletter of the American Malacological Union* 17 (2): 10.

129. Voight J. R. 1988. *Octopus Digueti* in Bahia Cholla, Sonora, Mexico. *Western Society of Malacology Annual Report (for 1987)* 20: 25–26.

130. Voight J. R. 1990. Population Biology of Octopus Digueti and the Morphology of American Tropical Octopods. University of Arizona. Dissertations PhD Thesis.

131. Váldez-Casillas C. 1996. Development and Testing of a Procedural Model for the Assessment of Human Wetland Interaction in the Tobari System of the Sonoran Coast Mexico. Tesis de Doctorado. Oregon State University. 163 p.

132. Watson M. C., Wayne R. F. 1991. A New Species of Suaeda (Chenopodiaceae) from Coastal Northwestern Sonora, Mexico. *Madroño* 38 (1): 30–36.

133. Watson M. K., Flessa K. W. 1979. Time-Averaged Molluscan Assemblages from Cholla Bay, Gulf of California. *Geological Society of America. Programs with Abstracts* 11 (3): 134.

134. Weeks J. R. 1986. The Growth and Water Relations of a Coastal Halophyte, *Salicornia Bigelovii*. PhD. Dissertation, Dept. of Molecular and Cellular Biology, University of Arizona.

135. Westervelt C. A. Jr. 1967. The Littoral Anomuran Decapod Crustacean Fauna of the Punta Peñasco-Bahía La Cholla Area in Sonora, Mexico. The University of Arizona. Ph.D. Dissertation. 159 pp

136. Williams, Austin B. 1986. Mud Shrimps, Upogebia, from the Eastern Pacific (*Thalassinoidea*, *Upogebiidae*). In. Memoirs of the San Diego Society of Natural History. Vol. 14, 1-60

137. Zamora H. A., Wilder B.T., Eastoe C.J., McIntosh J.C., Welker J., Flessa K.W. 2019. Evaluation of Groundwater Sources, Flow Paths, and Residence Time of the Gran Desierto Pozos, Sonora, Mexico. *Geosciences Journal* 9 (9): 378.

138. Zeh D. W. 1978. Community Structure and Factors Influencing the Distribution of Vascular Plants at Estero de La Cholla, Sonora, Mexico. In , XIII:28. Biological Studies in the Gulf of California. Vol. XIII (collected student papers from U. of Ariz. Marine Ecology Class), 28 pp

139. Zeh M. 1978. The Effects of Ray Foraging Disturbance on the Intertidal Infauna of Cholla Bay, Sonora, Mexico. In , XIII:22. Biological Studies in the Gulf of California.

140. Zerai D.B. 2007. Halophytes for Bioremediation of Salt Affected Lands. University of Arizona.Electronic Dissertation. PhD

**Table S3. Demographic characteristics of study participants**

| Location | Ejido Campodonico | Golfo de Santa Clara | Puerto Peñasco |
|---|---|---|---|
| *No. participants* | 17 | 6 | 25 |
| *Average age (± SE)* | 57.29±4.22 | 64±5.41 | 61.76±3 |
| Max age | 85 | 85 | 89 |
| Min age | 32 | 49 | 40 |
| *Main economic activity* | | | |
| Fishing | 10 | 0 | 8 |
| Aquaculture | 7 | 6 | 17 |
| *Average time residing in the community by main economic activity* | | | |
| Fishing | 43±3.4 | 53±3.6 | 47±9.8 |
| Aquaculture | 37±1.6 | 26±3.9 | — |
| *Employment status by main economic activity* | | | |
| Fishing | | | |
| Working | 6 | 3 | 10 |
| Retired | 3 | 3 | 7 |
| Aquaculture | | | |
| Working | 5 | — | 8 |
| Retired | 3 | — | 0 |
| *Participation in different fisheries by main economic activity* | | | |
| Fishing | | | |
| Small-scale | 7 | 10 | 3 |
| Subsistence | 0 | 7 | 3 |
| Sport fishing | 0 | 0 | 0 |
| Aquaculture | | | |
| Small-scale | 10 | 4 | — |
| Subsistence | 0 | 0 | — |
| Sport fishing | 0 | 0 | — |
| *Average years working by main economic activity* | | | |
| Fishing | 21±6 | 37.94±3 | 38.83±4 |
| Aquaculture | 22.1±4 | 18±3 | – |

**Figure S1. Output membership function. Divided into ten categories where 1 is the lowest abundance and 10 is the highest.**

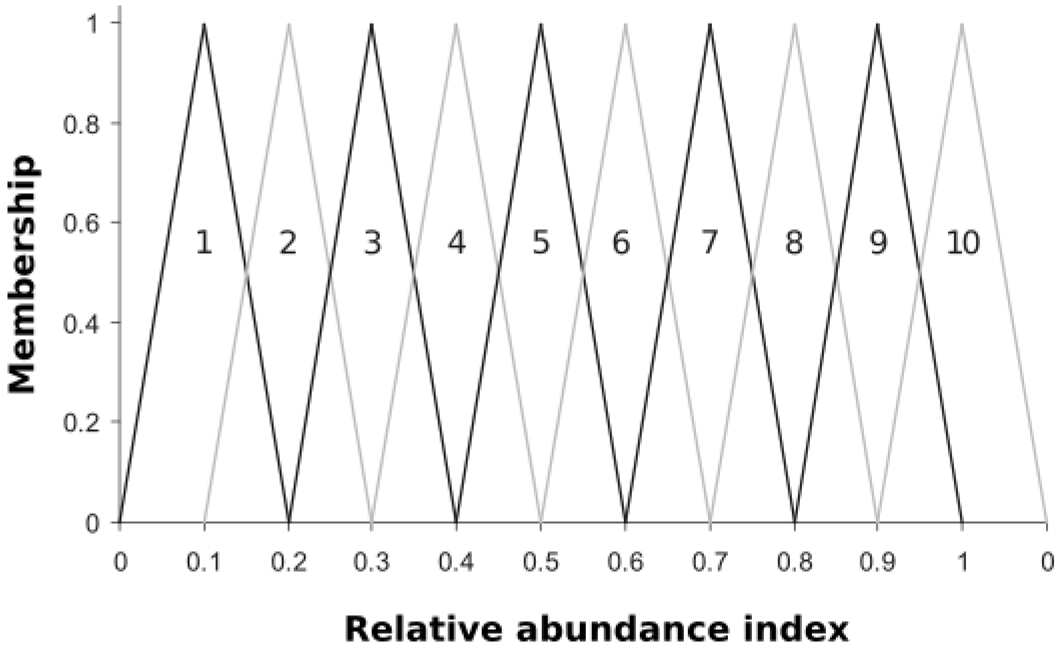

**Figure S2.** Uncorrected interview results. Proportion of fishers reporting high (dark grey), medium (medium grey), and low abundance (light gray) for each of 17 target species extracted in wetlands of the Northern Gulf of California.

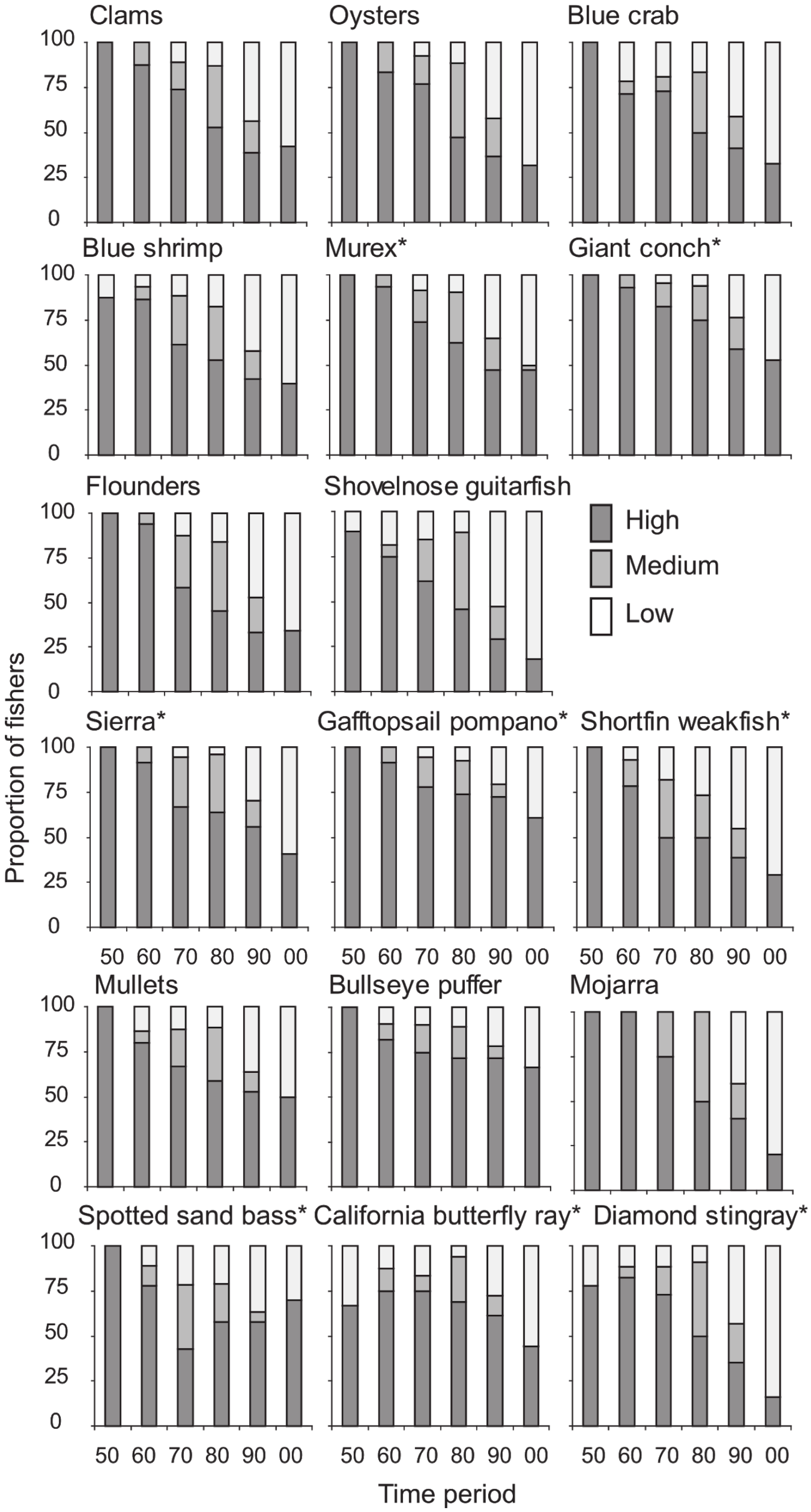

**Figure S3. Example of participatory drawing**

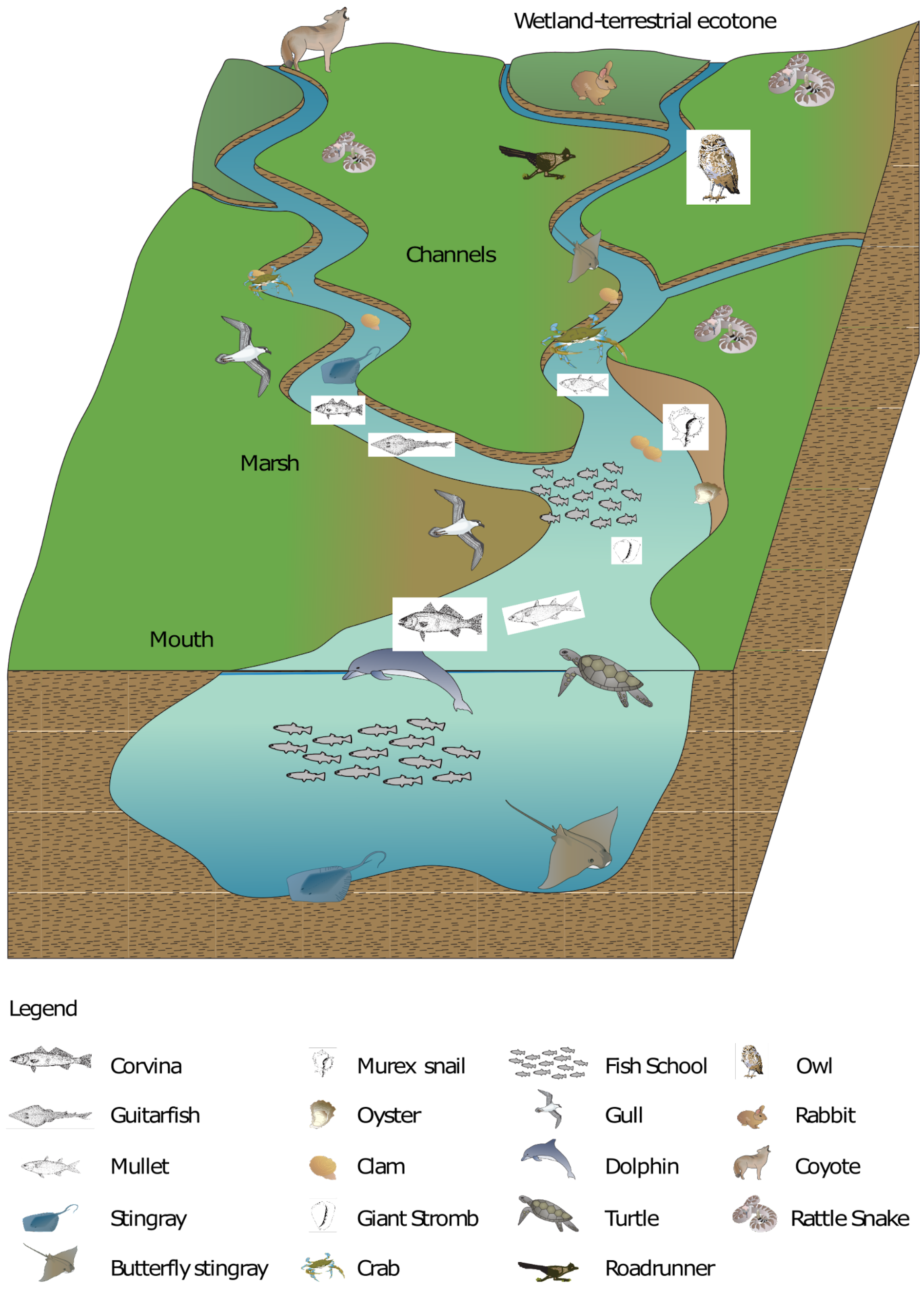

Wetland:La Cholla
Community:Puerto Peñasco
Date:05/16/08
h Survey number:05

**Figure S4. Historical ecology timetable of coastal wetlands in the Northern Gulf of California.**

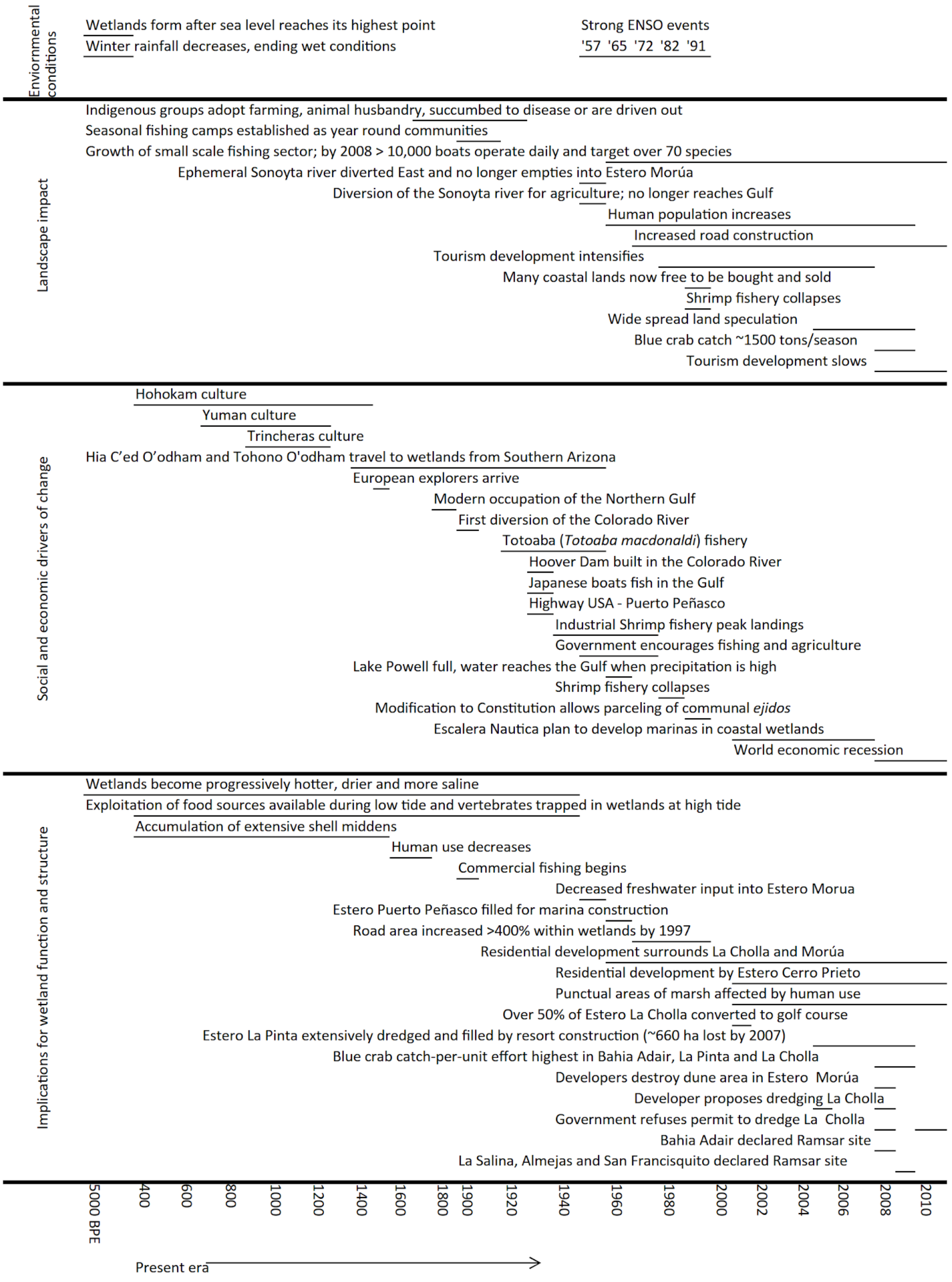
